# Non-cell-autonomous regulation of mTORC2 by Hedgehog signaling maintains lipid homeostasis

**DOI:** 10.1101/2024.05.06.592795

**Authors:** Kylie R. VanDerMolen, Martin A. Newman, Peter C. Breen, Laura A. Huff, Robert H. Dowen

## Abstract

Organisms must appropriately allocate energetic resources between essential cellular processes to maintain homeostasis and in turn, maximize fitness. The nutritional and homeostatic regulators of energy homeostasis have been studied in detail; however, how developmental signals might impinge on these pathways to govern cellular metabolism is poorly understood. Here, we identify a non-canonical role for Hedgehog (Hh), a classic regulator of development, in maintaining intestinal lipid homeostasis in *C. elegans*. We find that expression of two Hh ligands, GRD-3 and GRD-4, is controlled by the LIN-29/EGR transcription factor in the hypodermis, where the Hh secretion factor CHE-14/Dispatched also facilitates non-cell autonomous Hh signaling. We demonstrate, using *C. elegans* and mouse hepatocytes, that Hh metabolic regulation does not occur through the canonical Hh transcription factor, TRA-1/GLI, but rather through non-canonical signaling that engages mTOR Complex 2 (mTORC2) in the intestine. Hh mutants display impaired lipid homeostasis, including reduced lipoprotein synthesis and fat accumulation, decreased growth, and upregulation of autophagy factors, mimicking loss of mTORC2. Additionally, we found that Hh inhibits p38 MAPK signaling in parallel to mTORC2 activation and that both pathways act together to modulate of lipid homeostasis. Our findings show a non-canonical role for Hedgehog signaling in lipid metabolism via regulation of core homeostatic pathways and reveal a new mechanism by which developmental timing events govern metabolic decisions.

## INTRODUCTION

Proper allocation of resources between energetically expensive cellular processes is required for the survival of all organisms. Complex signaling networks govern metabolic commitments throughout the life of an animal to license growth and reproduction when environmental resources are available, thereby maximizing fitness of the species. Both developmental and environmental cues engage hubs of metabolic regulation to coordinate these energetic decisions, which maintain homeostasis at the cellular and organismal levels. While the core regulators of metabolism have been studied in detail, the mechanisms by which these regulatory pathways are tuned by developmental inputs remains poorly understood.

In *Caenorhabditis elegans*, maternal provisioning of lipids into the embryos is crucial for ensuring fitness of the progeny; however, this metabolic reallocation represents a metabolic trade-off that reduces maternal lifespan and dampens stress resilience^1–3^. These maternally provisioned lipids, which are stored in the intestine throughout development, are transported to the germline by triglyceride-rich low density-like lipoprotein (LDL-like) particles at the onset of adulthood via a process called vitellogenesis^4^. The temporal regulation of lipoprotein synthesis serves as a model to investigate how developmental signaling molecules engage metabolic pathways. Indeed, we have previously demonstrated that vitellogenesis is coordinated by a developmental timing program in the *C. elegans* hypodermis (*i.e.*, the skin) that couples to intestinal mTOR (<u>m</u>echanistic <u>T</u>arget <u>o</u>f Rapamycin) signaling, which in turn promotes lipoprotein synthesis^5^. The identity of the signaling molecules that mediate this inter-tissue crosstalk are unknown.

The *C. elegans* LIN-29 protein, a Krüppel-like zinc finger transcription factor that shares homology with the mammalian early growth response (EGR) proteins, is responsible for initiating several adult-specific developmental programs in the hypodermis, where it accumulates at the larval-to-adult transition^6–8^. Additionally, LIN-29 acts non-cell-autonomously to regulate intestinal lipoprotein synthesis by promoting the up-regulation of the vitellogenin genes (*vit-1-6*) at the larval-to-adult transition^5^. However, the role of LIN-29 in modulating intestinal metabolism is broader, as mutation of *lin-29* results in intestinal lipid storage defects; and consistently, ectopic over-expression of *lin-29* alters the expression of lipid metabolism genes and levels of specific fatty acids^5,9^. These effects on metabolism are presumed to be indirect, resulting from the misexpression of LIN-29 target genes in the hypodermis; however, the genes that are directly regulated by LIN-29 are unknown.

While the hypodermal-derived signaling molecules that engage intestinal metabolism are unknown, we have previously demonstrated that mTOR is one metabolic regulator that acts downstream of the hypodermal developmental timing program in the intestine to govern lipid homeostasis^5^. The evolutionarily conserved Ser/Thr protein kinase mTOR resides in two multiprotein complexes, mTOR Complex 1 (mTORC1) and mTOR Complex 2 (mTORC2), which together regulate several aspects of cellular metabolism and growth. Although the mechanisms of mTORC1 activation have been extensively studied, the upstream regulators of mTORC2 are poorly understood. In *C. elegans*, loss of the mTORC2-specific scaffolding protein RICT-1/Rictor results in impaired lipid homeostasis, reduced lipoprotein synthesis, slowed developmental rates, and a smaller body size^5,10,11^. Additionally, inappropriate induction of autophagy in the *rict-1* mutant confers a shorter lifespan^12,13^. Loss of SGK-1 (serum/glucocorticoid-regulated kinase 1), a downstream target of mTORC2, yields the same metabolic, growth, and aging phenotypes^5,11–15^. Additionally, it is well established that mTORC2 directly phosphorylates AKT in its hydrophobic motif to regulate growth and metabolism in mammalian cells^16–18^, which is likely conserved in *C. elegans*^19^. How intestinal mTORC2 signaling is engaged by the hypodermal developmental timing pathway to govern the activity of SGK-1 and AKT is not known.

Here, we find that Hedgehog (Hh) is the inter-tissue signaling molecule that couples developmental events in the hypodermis to metabolic regulation in the intestine. While Hh signaling is a well-characterized regulator of tissue patterning and morphogenesis in *Drosophila* and in vertebrates^20–22^, its role in governing lipid homeostasis is not well understood^23^. Canonical Hh signaling is initiated by binding of the mature Hh ligand to its receptor Patched, which results in de-repression of the seven-pass transmembrane protein Smo and activation of the Gli transcription factors^23^. We demonstrate that expression of two Hh-related ligands, GRD-3 and GRD-4, is controlled by LIN-29 in the *C. elegans* hypodermis. Dispatched/DISP, or CHE-14 in *C. elegans*, is required for Hh ligand secretion^24^, and consistently, *che-14* mutants display similar metabolic and growth phenotypes as the *grd-3; grd-4* double mutant, which includes decreased lipoprotein expression, impaired fat storage, and reduced body size. Importantly, we find that Hh signaling activates mTORC2 in both *C. elegans* and mouse hepatocytes independently of TRA-1/Gli, suggesting that Hh acts via a non-canonical pathway to govern mTORC2 activity and lipid homeostasis. In parallel, Hh represses the PMK-1/p38 MAPK pathway to promote lipid homeostasis and suppress p38-mediated transcriptional programs. Thus, Hedgehog signaling is a linchpin regulator of two homeostatic signaling pathways that is responsible for simultaneously committing metabolic resources to reproduction via mTORC2 signaling and restricting the metabolic flux towards metabolically expensive cellular processes (*i.e.*, stress and innate immunity pathways). Critically, investigation of the molecular mechanisms that underly this inter-tissue signaling event in *C. elegans* provides a robust system to identify new regulators of non-canonical Hh signaling that govern metabolic homeostasis.

## RESULTS

### LIN-29 regulates the expression of the Hedgehog-related ligands GRD-3 and GRD-4

Accumulation of the LIN-29/EGR transcription factor marks the final developmental event in the *C. elegans* hypodermis and is required for initiation of several adult-specific programs, including the synthesis of intestinal vitellogenin-containing lipoproteins^5–7^. We hypothesized that LIN-29 controls the transcription of an inter-tissue signaling factor that is produced in the hypodermis and is sensed by the intestine. To identify candidate signaling factors, we performed mRNA sequencing (mRNA-Seq) of transcripts from wild-type and *lin-29* mutant animals, finding 3,206 up-regulated and 2,752 down-regulated genes in the *lin-29* mutant (Figure S1A). We focused exclusively on the down-regulated genes, thereby assuming that LIN-29 regulates a positive regulator of intestinal metabolism, and compared these genes to a list of genes whose expression was restored in the *lin-29* mutant upon hypodermal, but not intestinal, rescue of *lin-29*. This resulted in the identification of 60 genes that were positively regulated by LIN-29 in the hypodermis (Figure S1A), which included three vitellogenin genes that we previously demonstrated were non-cell-autonomously regulated by *lin-29*^5^. Remarkably, this list contained only one candidate gene, *grd-3*, that was predicted to encode a secreted signaling molecule. GRD-3 is one of ∼60 Hedgehog-related proteins in *C. elegans* and contains a *groundhog* (*grd*) domain, which serves as the biologically active “Hedge” signaling component, but lacks the “Hog” domain responsible for autoproteolytically processing the bipartite Hedgehog proteins in *Drosophila*, vertebrates, and likely some *C. elegans* Hh-related proteins^25^. Due to the high functional redundancy of Hh-related ligands in *C. elegans*^25,26^, we examined the transcript levels of all Hh-related genes in the *lin-29* mutant and found that only two, *grd-3* and *grd-4*, were significantly down-regulated (Figure 1A). Notably, *grd-3* and *grd-4* are paralogues that share >70% nucleotide identity^26^, suggesting that they may function together to regulate Hh signaling downstream of LIN-29.

**Figure 1.**
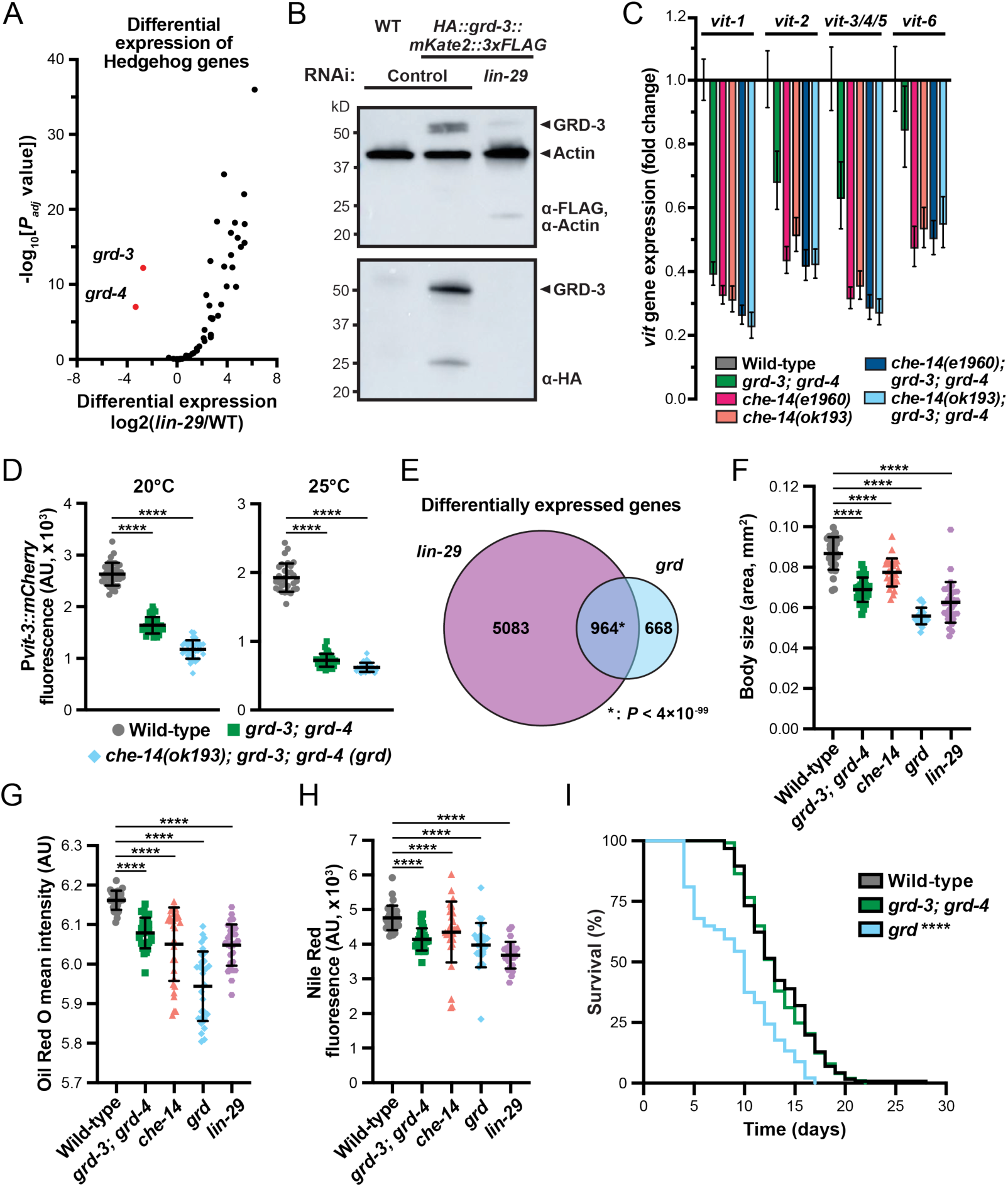
The LIN-29 transcription factor governs the expression of Hedgehog ligands that are required to maintain intestinal lipid homeostasis. (A) A volcano plot showing the differential expression of 58 Hh-related ligands in the *lin-29(n333)* mutant compared to wild-type (WT) reveals that *grd-3* and *grd-4* are strongly down-regulated. (B) A western blot using anti-FLAG and anti-actin (top) and anti-HA (bottom) of lysates from day 1 adult animals expressing HA::GRD-3::mKate2::3xFLAG protein at endogenous levels following *lin-29* RNAi. (C) Expression of the vitellogenin (*vit*) genes in day 1 adults reared at 20°C as determined by qPCR for the *grd-3(ok2778); grd-4(rhd134)* double mutant, as well as the indicated *che-14* single mutants and the *grd-3; grd-4; che-14* triple mutants. (D) Quantification of P*vit-3::mCherry* vitellogenesis reporter expression in day 1 adult wild-type or Hh mutant animals reared at 20°C (left) 25°C (right) (mean ± SD, ****, *P*<0.0001, one-way ANOVA). (E) A Venn diagram comparing the differentially expressed genes in the *lin-29(n333)* mutant to the *grd-3(ok2778); grd-4(rhd134); che-14(ok193)* (*grd*) triple mutant (hypergeometric *P* value reported). (F) Body size measurements and quantification of (G) Oil Red O and (H) Nile Red staining of day 1 adult wild-type, *lin-29(n333)*, and Hh mutant animals reared at 25°C. All data are reported as the mean ± SD (****, *P*<0.0001, ***, *P*<0.001, *, *P*<0.05, one-way ANOVA). (I) A longitudinal lifespan analysis of wild-type and the indicated Hh mutant animals cultured at 25°C (****, *P*<0.0001 between wild-type and *grd*, Log-rank test).

While *grd-3* transcript levels are dramatically down-regulated in the *lin-29* mutant, it’s unclear whether this results in an organism-wide loss of GRD-3 protein. Therefore, we inserted multiple epitope tags into the *grd-3* locus using CRISPR/Cas9, placing an HA tag following the signal sequence and a mKate2-3xFLAG tag at the C-terminus, which results in expression of a HA::GRD-3::mKate2::3xFLAG protein at endogenous levels. Importantly, insertion of these tags did not impair *grd-3* function (Figure S1B). Knockdown of *lin-29* by RNAi resulted in almost a complete loss of the GRD-3 protein, as revealed by western blotting with either anti-HA or anti-FLAG antibodies (Figure 1B). Intriguingly, we did detect a smaller band in the HA blot (∼25 kDa), which leaves open the possibility that GRD-3 might be cleaved in trans by a Hog domain-containing protein. Together, these data demonstrate that LIN-29 controls the levels two Hh-related proteins, which may, at least in part, underly the non-cell-autonomous effects conferred by loss of *lin-29*.

### The GRD-3 and GRD-4 ligands regulate intestinal lipid homeostasis

We reasoned that if GRD-3 and GRD-4 act redundantly to mediate the non-cell-autonomous effects of LIN-29, then loss of both genes should result in intestinal lipid metabolism defects that phenocopy the *lin-29* mutant. Thus, we first examined the mRNA levels of the vitellogenin (*vit*) genes in a *grd-3; grd-4* double mutant, finding that their loss reduced expression of all six *vit* genes (Figure 1C). Mutations in *che-14*, an orthologue of Dispatched/DISP, yielded mildly stronger defects in *vit* gene expression compared to the *grd-3/4* double mutant, but failed to dramatically enhance the *grd-3/4* mutations. Consistently, we found that expression of two vitellogenin transcriptional reporters, P*vit-2::GFP* and P*vit-3::mCherry*, were strongly decreased in the *grd-3; grd-4* double mutant and the *grd-3; grd-4; che-14* triple mutant, which we refer to as *grd* hereinafter (Figures 1D and S1C-D). This phenotype was slightly more severe at 25**°**C compared to 20**°**C. Since CHE-14/DISP would be predicted to facilitate secretion of numerous Hh-related proteins, it is not surprising that the *che-14* mutation more severely impairs *vit* gene expression relative to the *grd-3/4* double mutant; however, this also suggests that additional Hh-related ligands may work in concert with GRD-3/4 to regulate vitellogenesis.

Loss of *lin-29* resulted in misexpression of 5,958 genes, which can likely be attributed to both Hh-dependent and -independent effects. To assess the contribution of Hh signaling to *lin-29*-dependent gene expression, we performed mRNA-Seq and compared the differentially expressed genes (DEGs) between the *grd* and *lin-29* mutants. Remarkably, ∼60% of the genes misexpressed in *grd* are also differentially expressed in the *lin-29* mutant (Figure 1E), suggesting that GRD-3/4 regulates several, but not all, processes downstream of LIN-29. Consistent with having a broad role in regulating gene expression, the *lin-29* mutant displays both growth and metabolic phenotypes, including a smaller body size and reduced lipid storage^5^. Thus, we measured body size of the Hh mutants at adulthood and found that the *grd-3/4*, *che-14*, and *grd* mutant animals were all significantly smaller than wild-type animals (Figures 1F and S1E). Using Oil Red O or Nile Red to stain neutral lipids, we then assessed lipid levels in the Hh mutants. Consistent with our previous findings, mutations that disrupt *grd-3/4* or *che-14* impaired lipid accumulation in day 1 adults (Figures 1G-H and S1F-G), suggesting that proper Hh signaling from the hypodermis is required for accumulation of intestinal lipids, which could underly the smaller body size that we observed in the Hh mutants.

Dysregulation of lipid metabolism pathways can have profound effects on lifespan. For example, mutations in the insulin and mTOR signaling pathways, as well as dietary restriction interventions, can remodel lipid metabolism and autophagy pathways, resulting in an extended lifespan^27^. Given that impaired GRD-3/4 signaling reduced lipid levels, we hypothesized that the Hh mutants would display a dietary restriction-like increase in longevity. Surprisingly, the lifespan of the *grd-3/4* mutant was comparable to wild-type animals, while the *grd* triple mutant was significantly short lived (Figure 1I). These data demonstrate that CHE-14-dependent Hh signaling, independent of GRD-3/4, is required for longevity; however, this phenotype is unlikely to be driven by the reduced lipid levels since both the *grd-3/4* and the *grd* mutant display fat storage defects.

### Hypodermis-derived Hh signaling non-cell-autonomously regulates intestinal metabolism

LIN-29 acts in the hypodermis, or more specifically the hypodermal seam cells, to regulate intestinal vitellogenin expression (Figure S1A). Consistently, tissue-specific or single-cell mRNA-Seq experiments have shown that *grd-3*, *grd-4*, and *che-14* transcripts are enriched in hypodermal seam cells^28–30^. Thus, we hypothesized that LIN-29 may regulate the transcription of *grd-3* and *grd-4* in the seam cells, where they are then secreted by CHE-14/DISP to regulate lipid metabolism in a non-cell-autonomous manner. Upon inspection of animals expressing the HA::GRD-3::mKate2::3xFLAG protein, we observed strong mKate2 signal in the hypodermal seam cells and the surrounding hypodermal tissue, which was strongly diminished upon knockdown of *lin-29* by RNAi (Figure 2A). The mKate2 signal was distinct from body wall muscle cells, which are proximal to the hypodermal main body syncytium (Figure S2A). These data argue that LIN-29 regulates *grd-3* and *grd-4* expression in hypodermal seam cells.

**Figure 2.**
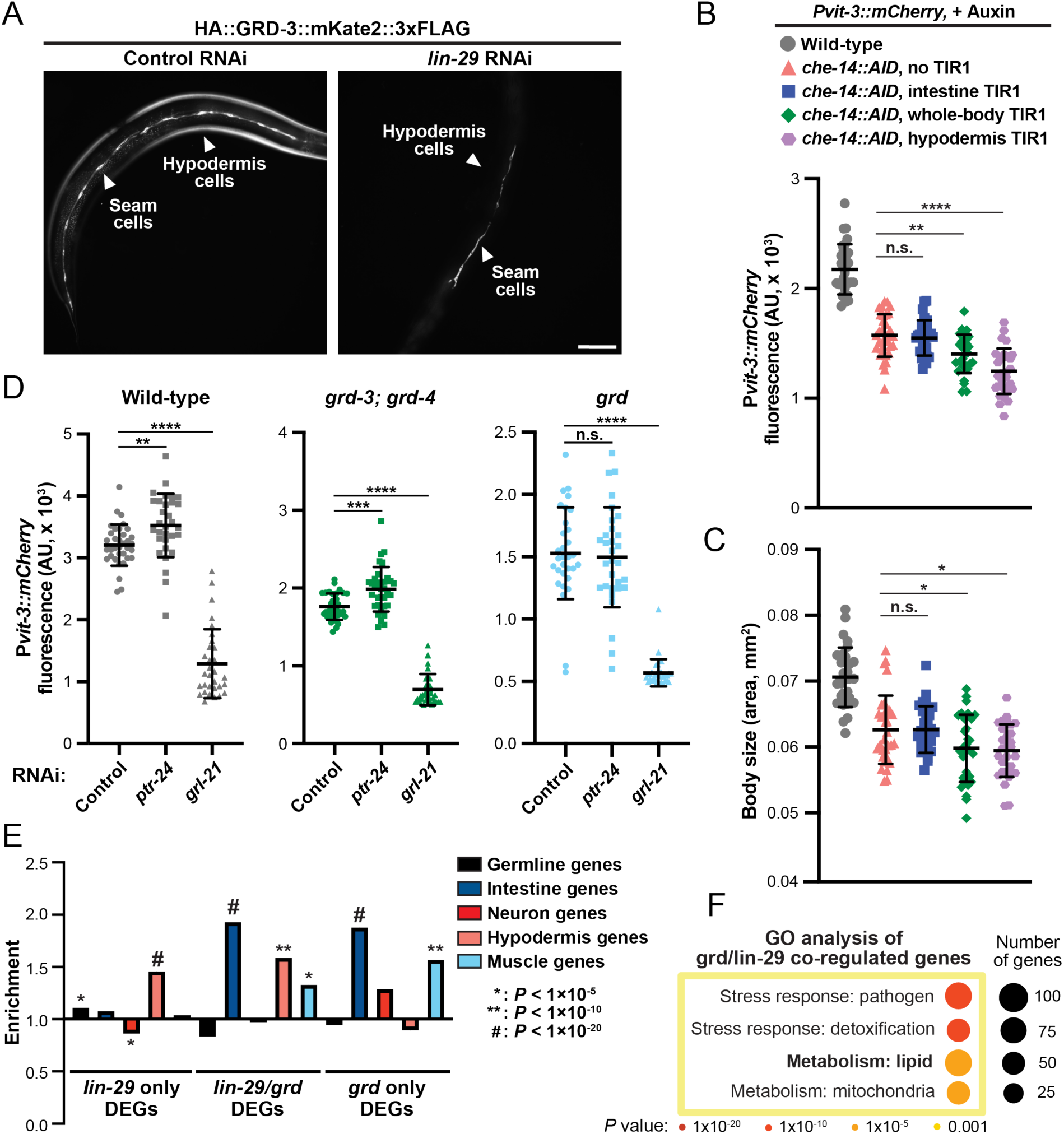
Hh ligands function non-cell-autonomously to control intestinal metabolism. (A) Representative flourescence images of animals expressing the GRD-3::mKate2 protein subjected to *lin-29* RNAi. GRD-3 is expressed in the hypodermal syncytia and seam cells (scale bar, 100μm). Quantification of (B) P*vit-3::mCherry* flourescence and (C) body size of *che-14::AID* animals cultured in the presence of 4 mM auxin at 20**°**C (mean ± SD, ****, *P*<0.0001, **, *P*<0.01, *, *P*<0.05, n.s., not significant, one-way ANOVA). The TIR1 protein is expressed from a single-copy transgene in a tissue-specific manner (intestinal, P*ges-1::TIR-1*; hypodermal, P*col-10::TIR-1*; whole body, P*eft-3::TIR-1*). (D) Quantification of P*vit-3::mCherry* reporter expression in the indicated strains following control, *ptr-24*, or *grl-21* RNAi (20**°**C, mean ± SD, ****, *P*<0.0001, ***, *P*<0.001, **, *P*<0.01, n.s., not significant, one-way ANOVA). (E) Enrichment (bars >1) or depletion (bars <1) of differentially expressed genes within the indicated tissues, indicating that *lin-29/grd* co-regulated genes are enriched in the intestine (relative to random chance). Hypergeometric *P* values are indicated. (F) A Gene Ontology analysis of the *lin-29/grd* co-regulated genes demonstrating that metabolic and detoxification pathways are under the control of LIN-29-Hh signaling (Fisher’s exact test *P* values are reported).

Secretion of Hh ligands is controlled by CHE-14/DISP. Thus, we predicted that CHE-14 would function in the hypodermis to control GRD-3/4 secretion and in turn, intestinal vitellogenin synthesis. To test this hypothesis, we engineered an auxin-inducible degron (AID) sequence into the *che-14* genomic locus using CRISPR/Cas9, which allows for tissue-specific depletion of AID-tagged proteins in the presence of auxin in cells expressing the TIR1 ubiquitin ligase. Since it was not clear whether the N- or C-termini could be accessed by TIR1, we introduced the AID tag into an internal loop of the multiple-pass transmembrane CHE-14 protein and expressed TIR1 under the control of either an intestinal, hypodermal, or ubiquitous promoter. Importantly, the AID tag partially reduced CHE-14 function compared to wild-type animals (Figure 2B-C). Therefore, we compared auxin-dependent depletion of CHE-14 in each tissue to the CHE-14::AID strain lacking TIR1, finding that ubiquitous and hypodermal depletion of CHE-14, but not intestinal depletion, further impaired P*vit-3::mCherry* reporter expression and reduced body size (Figures 2B-C and S2B).

Given that CHE-14 acts in the hypodermis to facilitate secretion of Hh-related ligands and that *che-14* mutations more severely impair *vit* gene expression compared to the *grd-3/4* mutant, we speculated that additional Hh-related proteins may be produced in the hypodermis that control intestinal lipoprotein synthesis. It was previously shown that another Hh-related protein, GRL-21, is transcriptionally regulated in the hypodermis and non-cell-autonomously promotes intestinal lipid accumulation by inhibiting PTR-24, a Patched-related receptor^31^. We reasoned that GRD-3/4 may work cooperatively with GRL-21 following CHE-14-dependent secretion from the hypodermis to regulate intestinal lipoprotein expression by inhibiting PTR-24. Indeed, knockdown of *grl-21* by RNAi dramatically reduced P*vit-3::mCherry* reporter expression in wild-type, *grd-3/4* double mutant, and *grd* triple mutant animals (Figure 2D). However, knockdown of *ptr-24* by RNAi or intestine-specific depletion of PTR-24::AID only had a modest effect on restoring P*vit-3::mCherry* expression in a *grd-3/4* mutant background (Figures 2D and S3A), suggesting that PTR-24 likely does not negatively regulate vitellogenesis. Thus, it is possible that control of intestinal lipoprotein production by GRD-3/4 and GRL-21 occurs through two distinct mechanisms that bind different Patched or Patched-related receptors, of which there are 26 in *C. elegans*^26^.

Our data suggest that Hh-related proteins are secreted from the hypodermis and act on distal tissues to regulate gene expression. Given that impaired Hh signaling resulted in strong down-regulation of the *vit* genes, we hypothesized that GRD-3/4 acts on intestinal cells to exert transcriptional control over various aspects of intestinal lipid metabolism that is required for the accumulation of lipids and synthesis of lipoproteins. Indeed, we found that *lin-29*/*grd*-coregulated genes, as well as *grd*-specific genes, were strongly enriched for intestinal expression (Figure 2E), indicating that the *lin-29* and *grd* mutations broadly alter intestinal gene expression, which is consistent with their pleiotropic effects. Using gene ontology (GO) enrichment^32^, we found that *lin-29*/*grd*-coregulated genes were enriched for functions in stress responses (immunity and detoxification) and lipid and mitochondria metabolism (Figure 2F). Together, our data argue that LIN-29 regulates the expression of Hh-related ligands, which are secreted from the hypodermis by CHE-14, and act on intestinal targets to govern lipid metabolism pathways.

### Hedgehog engages mTORC2 signaling to regulate metabolic homeostasis

We have previously demonstrated that the vitellogenesis defects in the *lin-29* mutant can be partially suppressed by hyper-activation of the mTORC2 signaling pathway^5^, suggesting that LIN-29-dependent signals may directly regulate mTORC2. Moreover, two direct targets of mTORC2, AKT-1 and SGK-1, are required for proper vitellogenin expression and growth^5,10,11,19^. Thus, we reasoned that hypodermal Hh-related ligands may act on receptors in the intestine to regulate mTORC2 via a non-canonical Hh signaling pathway. Despite the significant redundancy among the Patched and Patched-related receptors in *C. elegans*^26^, we sought out to identify these receptors. Using tissue-specific RNA-seq data as a guide^29^, we prioritized receptors with elevated expression in the intestine (*i.e.*, PTC-1, PTC-3, PTR-4, PTR-21, PTR-24) and performed intestine-specific RNAi to assess their role in vitellogenesis. Knockdown of *ptr-21* yielded a detectable decrease in *Pvit-3::mCherry* expression, which we recapitulated using four different *ptr-21* mutant alleles (Figures S3B-C). The modest defects produced by loss of *ptr-21* suggests that additional intestinal receptors are likely acting redundantly.

To more directly test whether Hedgehog signaling activates mTORC2, we chose to employ AML12 mouse hepatocytes, which express (at most) two Patched receptors, are responsive to the Hh pathway activator SAG (Smoothened Agonist)^33^, and lack primary cilia, making them an ideal model to investigate non-canonical Hh signaling. Moreover, mTORC2 regulatory mechanisms identified in *C. elegans* are conserved in AML12 cells^13^. Upon treatment of AML12 hepatocytes with SAG, we observed phosphorylation of Akt at Ser473, the site of mTORC2 regulation within the hydrophobic motif^16^, and Thr308 within the activation loop (Figure 3A). Furthermore, we observed SAG-induced phosphorylation of the FOXO transcription factor, a major downstream target of both Akt and SGK1. This phosphorylation event occurred rapidly in response to SAG and did not correlate with a significant of accumulation of Gli (Figure 3A). Next, we assessed whether *Rictor* was required for SAG-induced phosphorylation of Akt using shRNA knockdown, finding that depletion of Rictor abrogated SAG-induced phosphorylation of Akt^S473^ (Figure 3B). Together, these data suggest that Hedgehog signaling controls mTORC2 activity in hepatocytes.

**Figure 3.**
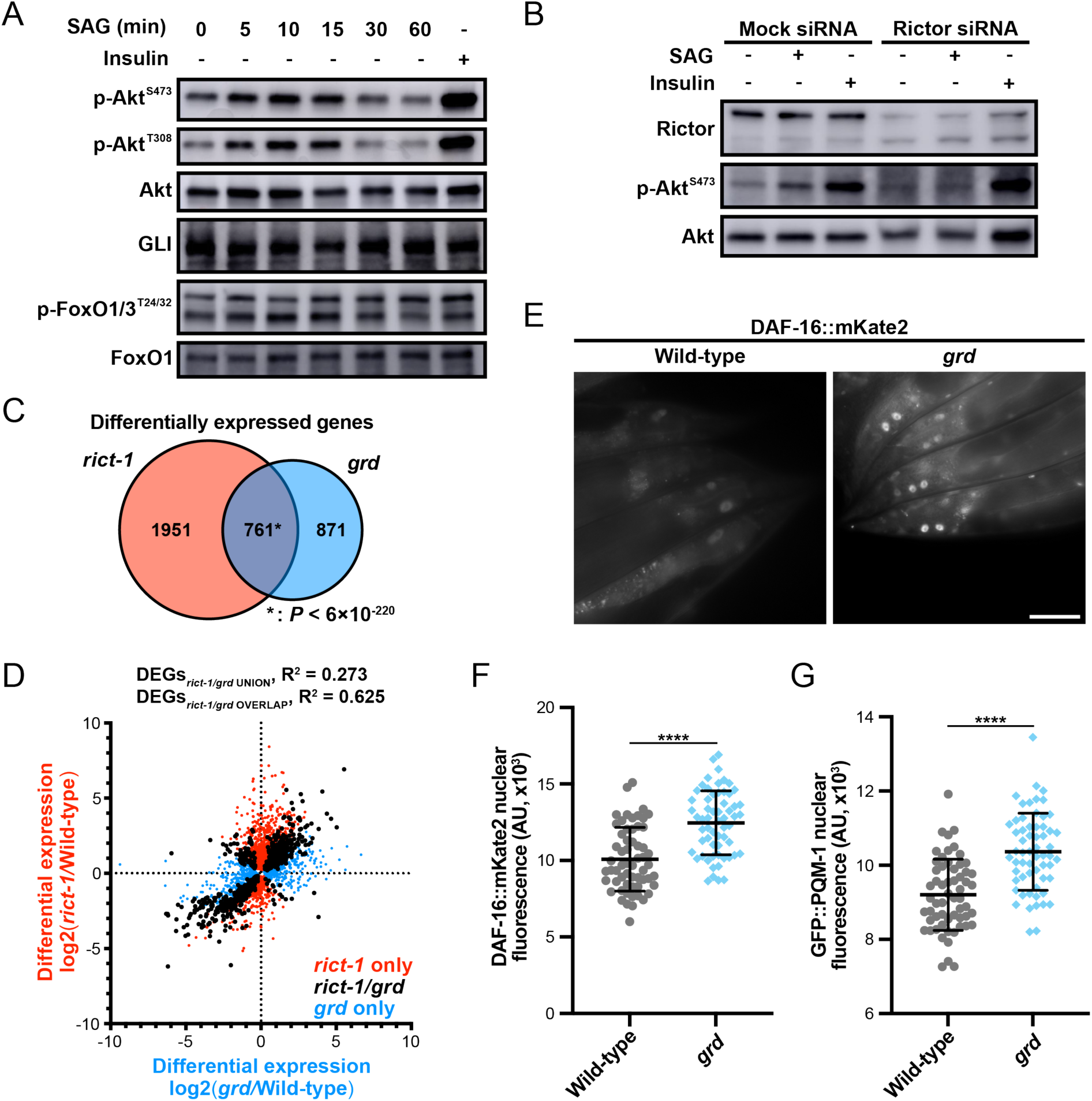
Hh stimulates mTORC2 signaling. (A) A western blot analysis of lysates from AML12 mouse hepatocytes showing that Smoothened Agonist (SAG) stimulates phosphorylation of Akt at Ser473, the mTORC2 regulatory site, and secondary Thr308 phosphorylation site without altering GLI1 levels. SAG also stimulates FoxO phosphorylation. Insulin treatment is a positive control for Akt phosphorylaton (B) SAG-induced phosphorylation of Akt Ser473 requires Rictor in AML12 hepatocytes. Cells were treated with either mock or Rictor siRNAs for 48 hours prior to vehicle, 0.5 μM SAG treatment (15 min), or 100 nM insulin treatment (30 min). (C) A Venn diagram comparing the differentially expressed genes in the *rict-1(mg360)* mutant to the *grd* triple mutant with the hypergeometric *P* value reported. (D) A scatter plot showing the differential expression values for the *rict-1(mg360)* mutant plotted against the *grd* triple mutant with the *rict-1*-specific DEGs shown in red, the *grd*-specifc DEGs in blue, and the *rict-1/grd* co-regulated genes in black. A linear regression analysis was performed on the union and intersection of the *rict-1* and *grd* datasets with the individual R^2^ values reported. (E) Representative images (scale bar, 50 μm) and (F) quantification of DAF-16::mKate2 nuclear flourescence in day 1 adult wild-type and *grd* mutant animals reared at 20**°**C (mean ± SD, ****, P<0.0001, T-test). (G) Quantification of GFP::PQM-1 nuclear flourescence revealed an increase in nucear accumulation of PQM-1 in the *grd* mutant (****, P<0.0001, T-test).

In *C. elegans*, *rict-1/Rictor* is required for initiation of lipoprotein synthesis at adulthood^5^, which also requires *grd-3/4*. If Hh regulates mTORC2 activity in *C. elegans*, as observed in hepatocytes, we would expect that disruption of either Hh or mTORC2 would result in misexpression of the same genes, including the *vit* genes. Therefore, we performed mRNA-Seq on the *rict-1* mutant and compared the DEGs to those genes differentially expressed in the *grd* mutant. This analysis revealed a striking overlap between these datasets with ∼47% of the *grd*-dependent genes also mis-expressed in the *rict-1* mutant (Figure 3C). Furthermore, the 761 co-regulated genes, which were enriched for intestinal genes, were mis-expressed to similar levels in both mutants (Figures 3D and S4A). Consistent with these observations, two different *sgk-1* gain-of-function mutations [unpublished work] partially suppressed the vitellogenesis and small body size of the *grd-3/4* mutant (Figures S4B-E). Together, these results suggest that GRD-3/4 acts in the same pathway as mTORC2.

To gain insight into the transcription factors (TFs) that regulate the *rict-1*/*grd* co-regulated genes, we assessed nuclear localization of metabolic transcription factors downstream of mTORC2, DAF-16 and PQM-1, using fluorescent reporters. Loss of *grd* prompted a robust accumulation of DAF-16 and PQM-1 in the nucleus (Figure 3F). Nuclear accumulation of DAF-16 resulting from impaired Hh signaling in *C. elegans* is consistent with SAG-induced phosphorylation of FOXO in hepatocytes (Figure 3A). These data are consistent with a Hedgehog-mTORC2 signaling axis to regulate metabolic transcription.

### Impaired Hedgehog signaling stimulates autophagy

The mTORC2-SGK-1 pathway is known to impact some aspects of autophagy, including negative regulation of the VDAC1 ion channel and inhibition of mitochondrial permeability, which is required for autophagy-dependent lifespan extension^13^. Consistently, *rict-1* and *sgk-1* mutants show an increase in the number of autophagic vesicles, up-regulation of master transcriptional regulator of autophagy HLH-30/TFEB^34^, and a shortened lifespan that can be restored by reducing autophagic programs^13^. We hypothesized that Hedgehog modulates autophagy by engaging intestinal mTORC2 signaling in *C. elegans*. To test whether autophagy is elevated upon loss of Hh signaling, we measured the number of autophagosomes marked by GFP::LGG-1, the *C. elegans* orthologue of Atg8. Indeed, we found an increase in number of GFP::LGG-1 puncta in the proximal intestine, as well as an increase in GFP::LGG-1 protein levels and proteolytic processing, in the *grd* mutants (Figures 4A-C), suggesting that Hh signaling restricts autophagy similar to that of mTORC2-SGK-1 signaling^13^. Using our mRNA-Seq data, we then examined mRNA levels of *hlh-30*, finding that *hlh-30* transcripts were increased in the *lin-29, grd*, and *rict-1* mutants (Figure 4D). Using the translational reporter HLH-30::GFP, we next inspected localization of HLH-30 in the *grd* and *rict-1* mutant backgrounds. Consistent with *hlh-30* transcript levels, HLH-30::GFP nuclear localization was increased in response to impaired Hh or mTORC2 signaling (Figures 4E-F). Given that elevated autophagy restricts lifespan in the *rict-1* mutant^13^, we performed *hlh-30* RNAi on *rict-1* and *grd* mutants and found that knockdown of *hlh-30* extended the lifespan of both mTORC2 and Hh mutants (Figure 4G). Together, our data suggest that Hh signaling, like mTORC2, represses autophagic programs, at least partially through the action of HLH-30, to govern organismal homeostasis.

**Figure 4.**
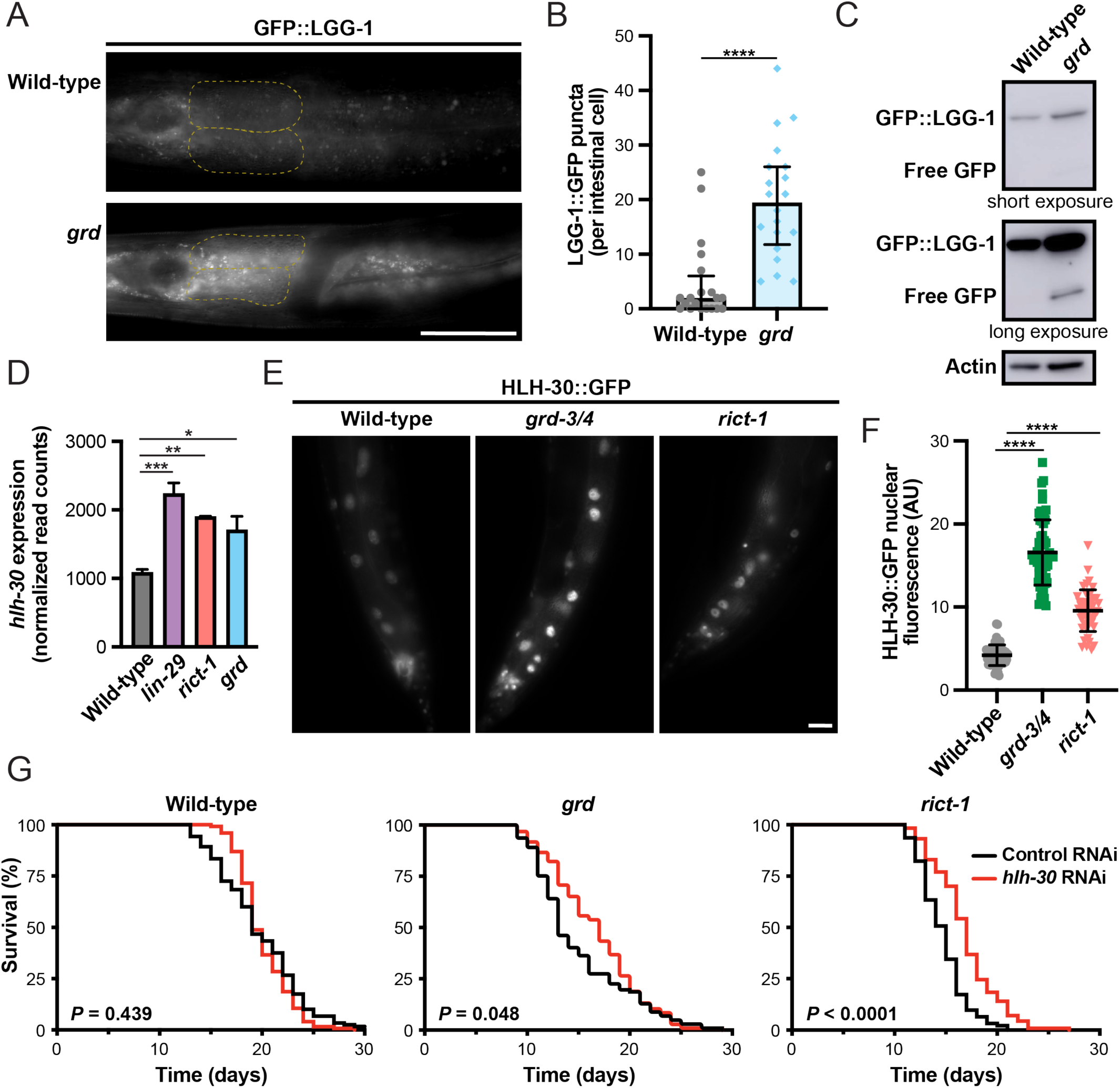
Impaired Hh signaling induces autophagy programs. (A) Representative images (scale bar, 50 μm) and (B) measurement of number GFP::LGG-1 puncta per intestinal cell (median with interquartile range, ****, *P*<0.0001, T-test) in the two proximal-most intestinal cells (outlined in yellow-dashed lines in A) in day 1 adult wild-type and *grd-3(ok2778); grd-4(rhd134); che-14(ok193)* mutant animals. (C) A western blot analysis of lysates from day 1 adult wild-type or *grd* mutant animals expressing GFP::LGG-1 using anti-GFP (top and middle) and anti-actin (bottom) antibodies. (D) Normalized *hlh-30* expression levels (RPKM, reads per kilobase of transcript per million mapped reads) in wild-type, *lin-29(n333)*, *rict-1(mg360)*, and *grd* mutant animals (mean ± SEM, ****, *P*<0.0001, ***, *P*<0.001, *, *P*<0.01, one-way ANOVA). (E) Representative images (scale bar, 20 μm) and (F) quantification of HLH-30::GFP nuclear flourescence (mean ± SD, ****, *P*<0.0001, one-way ANOVA) in day 1 adult wild-type, *grd-3(ok2778); grd-4(rhd134)*, and *rict-1(mg360)* animals cultured at 25**°**C. (G) Longitudinal lifespans of wild-type, *grd*, and *rict-1(mg360)* animals reared at 20**°**C and subjected to either control or *hlh-30* RNAi. The Log-rank test *P* values are displayed.

### Hedgehog functions independently of TRA-1/Gli to govern intestinal gene expression

In vertebrates, canonical Hedgehog signaling alters transcription of target genes through accumulation of zinc finger transcription factor Gli^20,35^. The *C. elegans* orthologue of Gli, TRA-1, is a transcriptional regulator of sexual cell fate decisions, which is mediated in part through its repression of the male-specific transcription factor MAB-3^36,37^. It is not yet clear whether TRA-1 acts as the transcriptional effector of Hh signaling in *C. elegans.* Since we observed little Gli accumulation in hepatocytes in response to SAG, we predicted that Hh signaling is unlikely to govern metabolic pathways via TRA-1 in *C. elegans*. However, knockdown of *tra-1* in the intestine using a tissue-specific RNAi strain resulted in a stark reduction in P*vit-3::mCherry* expression (Figure 5A). To further assess the role of *tra-1* in regulation of intestinal metabolism, we performed mRNA-Seq on animals subjected to intestine-specific knockdown of *tra-1*. Upon inspection of the *rict-1*/*grd* co-regulated genes, we found that while the *lin-29, rict-1,* and *grd* mutations confer a very similar pattern of differential expression, *tra-1* intestine-specific knockdown resulted in a markedly different pattern, suggesting that TRA-1 is not a transcriptional effector of the LIN-29/Hh/mTORC2 signaling axis (Figure 5B). Moreover, loss of intestinal *tra-1* failed to up-regulate the autophagy factor *hlh-30*/*TFEB*; however, *mab-3* was strongly up-regulated, which was unique to *tra-1* knockdown (Figure 5C). These data argue that MAB-3, a male-specific repressor of vitellogenesis, is suppressing *vit* gene expression following loss of *tra-1*^37,38^. Consistent with this hypothesis, intestine-specific knockdown of *tra-1* further enhances the defects in *vit* gene expression conferred by a null mutation in *sgk-1* (Figure 5D), suggesting that mTORC2 and TRA-1 act in distinct pathways to regulate vitellogenesis. These data suggest that Hedgehog acts through mTORC2, and not the TRA-1 sex determination pathway, to govern intestinal homeostasis.

**Figure 5.**
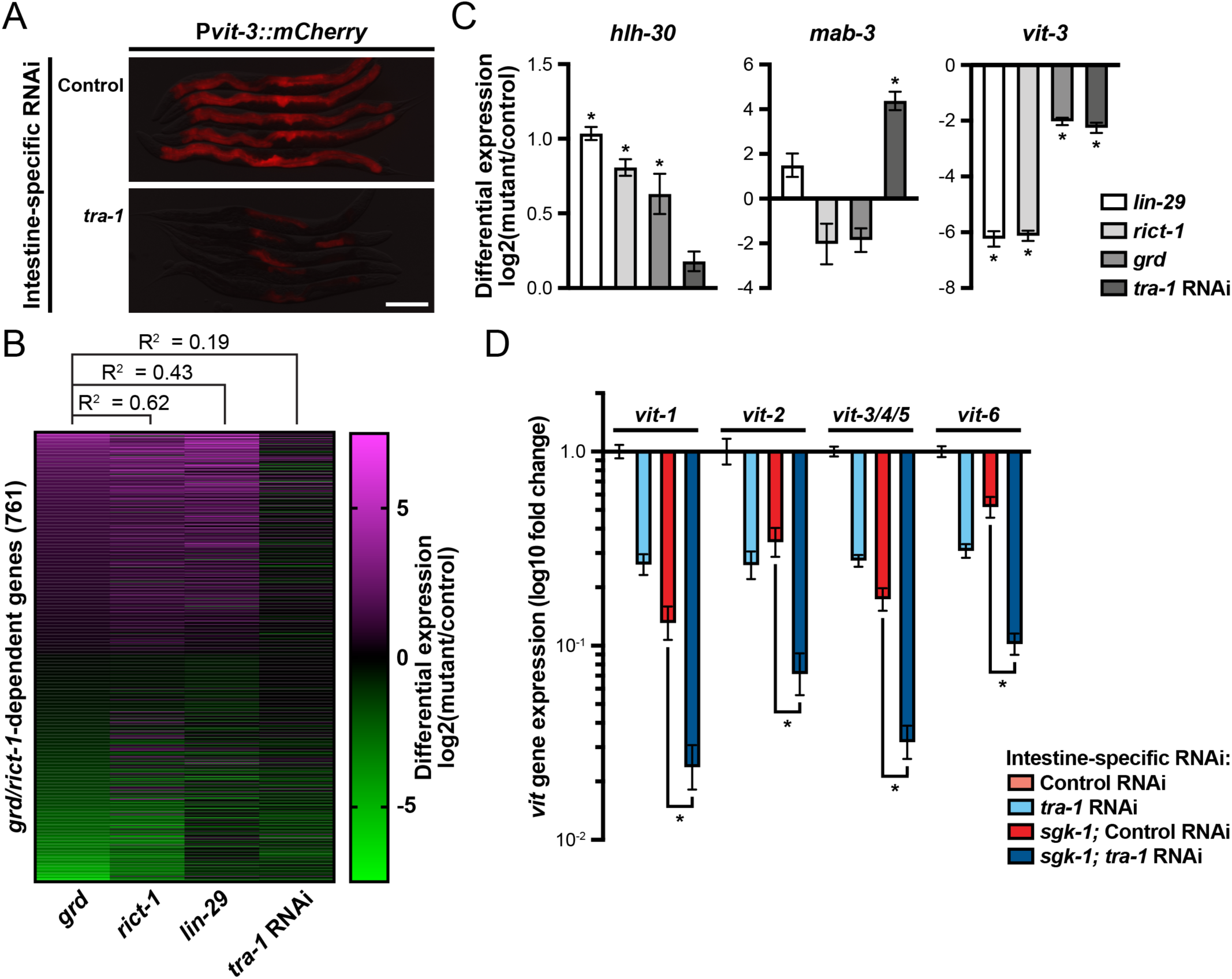
Hh signaling functions independently of the *C. elegans* GLI orthologue TRA-1. (A) Representative P*vit-3::mCherry* flourescence images of an intestine-specific RNAi strain subjected to control or *tra-1* RNAi (day 1 adults at 20**°**C; scale bar, 200 μm). (B) A heatmap of differential gene expression values derived from mRNA-Seq data for the 761 *rict-1/grd* co-regulated genes for the *grd*, *rict-1(mg360)*, and *lin-29(n333)* mutants (values relative to wild-type), as well as animals subjected to intestine-specific *tra-1* RNAi (values relative to control RNAi). The rows are ordered from the most up-regulated gene to most down-regulated gene in the *grd* sample and the R^2^ values are reported for each sample comparison relative to the *grd* sample. (C) Differential expression values (log2 fold change) for *hlh-30*, *mab-3*, and *vit-3* in the indicated mutant strains (*, P_adj_<0.01, DESeq2). (D) A qPCR analysis of vitellogenin (*vit*) gene expression in day 1 adult wild-type or *sgk-1(ok538)* animals subjected to control or *tra-1* intestine-specific RNAi at 20**°**C (*, P<0.01, T-test).

### Hedgehog signaling represses the p38 MAPK pathway

The signaling factors and transcriptional effectors downstream of non-canonical, Gli-independent Hedgehog signaling responsible for maintaining cellular homeostasis are largely unknown^23^. It is possible that the Hh-mTORC2 signaling axis might together regulate a homeostatic pathway. To investigate this possibility, we queried whether the *rict-1/grd* co-regulated genes are known to be regulated by any additional signaling pathways or transcription factors using the WormExp tool^39^. We found a strong enrichment for genes regulated by p38 MAPK pathway (Figure 6A), which is comprised of the NSY-1/SEK-1/PMK-1 kinase cascade under control of the conserved TIR-1/SARM1 Toll/interleukin-1 receptor (TIR) domain-containing protein. In *C. elegans*, activation of the p38 pathway induces innate immunity and oxidative stress responses while simultaneously restricting developmental rate^40–42^, possibly by balancing the metabolic commitment between these energetically expensive processes. Thus, we measured lipid levels in the *grd* and *rict-1* mutants following inactivation of the p38 pathway with a *tir-1* mutation. Loss of *tir-1* partially suppressed the lipid storage defects displayed by the Hh and mTORC2 mutants (Figure 6B), suggesting that p38 governs lipid pathways downstream or in parallel to the Hh-mTORC2 signaling axis. To assess whether p38 signaling acts broadly to govern mTORC2- and Hh-dependent transcriptional programs, we generated mRNA-Seq datasets for *tir-1; rict-1* and *tir-1; grd* mutants and calculated the differential expression levels for all 1,632 *grd*-dependent genes. Loss of *tir-1* dramatically suppressed the expression defects of those genes down-regulated, but not up-regulated, in the *grd* and *rict-1* mutants (Figures 6C-D). Together, these data suggest that p38 MAPK may play a role in lipid metabolic regulation through crosstalk with Hh and/or mTORC2 pathways.

**Figure 6.**
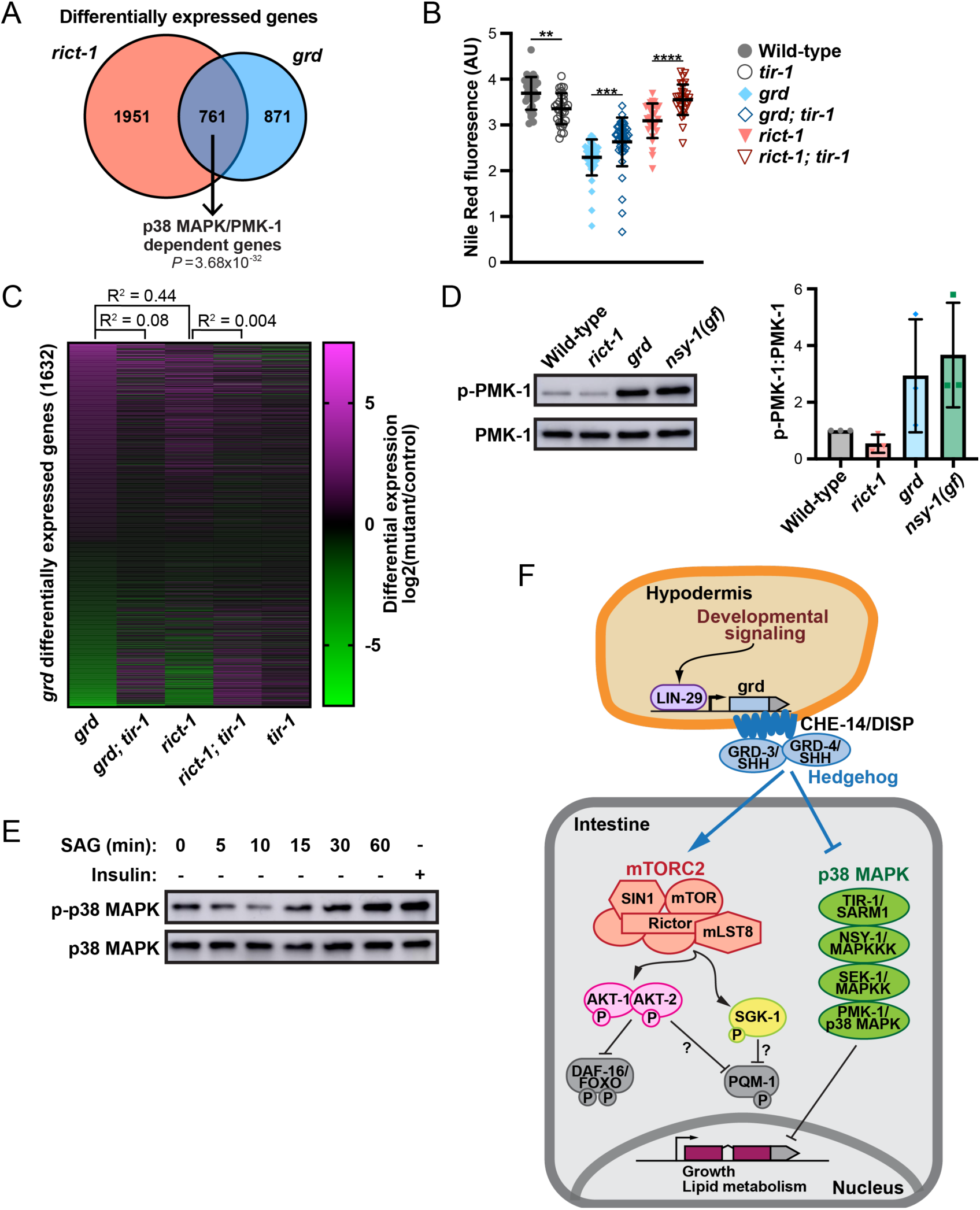
Hh inhibits p38 signaling to govern intestinal homeostasis. (A) A Venn diagram showing that the *rict-1/grd* co-regulated genes are enriched for *pmk-1*-dependent genes (hypergeometric *P* value reported^66^). (B) Quantification of lipid levels using Nile Red staining of the indicated wild-type and mutant strains. Data are representated as the mean ± SD (****, *P*<0.0001, ***, *P*<0.001, **, *P*<0.01, one-way ANOVA). (C) A heatmap of the 1,632 differentially expressed genes in *grd* mutants showing differential expression values (log2 fold change relative to wild-type) for the indicated mutant strains. Each row represents a gene organized from most up-regulated to most down-regulated in *grd* and R^2^ values are shown to compare *grd* to *grd; tir-1* and *rict-1*, as well as *rict-1* to *rict-1; tir-1*. Western blot analyses of phospho-p38 levels for (D) *C. elegans* lysates prepared from wild-type, *rict-1(mg360)*, *grd*, or *nsy-1(ums8)* mutants (*ums8* is a gain-of-function allele) and (E) AML12 hepatocytes treated with SAG for increasing amounts of time. The pPMK-1/PMK-1 ratio is plotted for the *C. elegans* data. (F) A model of how Hh governs intestinal metabolism through the dual regulation of mTORC2 and p38 signaling in *C. elegans*.

We reasoned that reduced Hh/mTORC2 signaling may hyperactivate p38 signaling, resulting in dysregulation of transcriptional programs and altered organismal homeostasis. To test this hypothesis, we examined the phosphorylation state of PMK-1, finding that impaired Hh signaling resulted in hyper-phosphorylation of PMK-1 in *C. elegans* (Figure 6E). Consistently, activation of Hh signaling using SAG reduced p38 phosphorylation in hepatocytes (Figure 6F). Surprisingly, loss of *rict-1* did not result in PMK-1 hyperphosphorylation (Figure 6E), suggesting that Hh signaling may repress p38 signaling independently of mTORC2. Together, our data suggest that Hh simultaneously activates mTORC2 while repressing p38 signaling to coordinate lipid homeostasis (Figure 6G).

## DISCUSSION

Here, we demonstrate that Hedgehog is a linchpin regulator of lipid metabolism and growth during development. The developmental timing factor, LIN-29, regulates the transcription of the Hh ligands GRD-3 and GRD-4, which engage downstream homeostatic signaling. Hedgehog non-cell-autonomously activates mTORC2 signaling while repressing the p38 pathway; however, both pathways impinge on a common set of genes to maintain intestinal homeostasis in *C. elegans* (Figure 6G). Importantly, this Hh signaling axis is conserved, as we find that Smo agonist stimulates mTORC2 activity and inhibits p38 signaling in mouse hepatocytes. Intriguingly, a Smo orthologue has not yet been identified in *C. elegans* using comparative genomic approaches; however, a functional Smo equivalent may have diverged significantly to evolve alongside of the ∼60 Hh ligands that are present in *C. elegans*. Nonetheless, our demonstration that the *C. elegans* Hh-mTORC2 signaling axis is conserved in mammalian cells is, to our knowledge, the first evidence supporting a role for non-canonical Hh signaling in metabolic regulation that is conserved from nematodes and mammals.

We extensively searched for the Patched or Patched-related receptor(s) that, like *grd-3* and *grd-4*, are required for proper vitellogenin expression in *C. elegans*; however, we were unable to identify a single receptor that carries out this function. It is possible that multiple Patched, Patched-related, or LRP (LDL-receptor-related protein^43^) receptors work redundantly to coordinate the activity of GRD-3 and GRD-4. Alternatively, additional hypodermis-derived Hh ligands may work in tandem on a shared suite of intestinal receptors to activate or repress various aspects of intestinal lipid metabolism. In support of this hypothesis, ectopic over-expression of *lin-29* induces transcription of the *wrt-6*, *grl-14* and *grd-11* Hh ligands, which repress the expression of some intestinal lipid metabolism genes^9^. Thus, loss of *grd-3* and *grd-4* might sensitize the intestine to other Hh ligands, or possibly other endocrine factors, that contribute to the lipid metabolism phenotypes that we observed in the Hh mutants. In contrast, activation of mTORC2 by Smo in hepatocytes, a system that lacks the redundancy seen in *C. elegans*, suggests that Hh signaling may directly engage mTORC2 rather than acting through a different endocrine or paracrine factor.

Non-canonical Hh signaling is known to regulate various aspects of lipid metabolism in *C. elegans*, including nutrient-dependent lipid accumulation by GRL-21/PTR-24^31^, regulation of cholesterol transport by PTC-3^44^, and repression of fatty acid desaturase gene expression by PTR-23^45^; however, it is unknown whether Hh signaling engages mTOR signaling in these contexts. Interestingly, GRD-1/PTR-11 represses growth rate upon inactivation of mTORC2 signaling^46^, yet the mechanism by which Hh coordinates growth in the absence of mTORC2 is unknown. Some of the most compelling evidence for a Hh-mTOR signaling axis in developmental regulation is in *D. melanogaster* eye, where Hh instructs spatial activation of mTORC1 and subsequent cell divisions^47^. Notably, this developmental event is coordinated at the transcriptional level by the Gli orthologue Cubitus interruptus (Ci). We found that the *C. elegans* Gli orthologue TRA-1 and GRD-3/4 regulate a distinct set of genes, suggesting Gli is unlikely to be regulating mTORC2 at the transcriptional level. Consistently, GLI1 levels are not altered upon acute SAG treatment of AML12 hepatocytes. In contrast, hepatocyte-specific ablation of Smo in mice causes a chronic imbalance in GLI1, GLI2, and GLI3 transcriptional activity, resulting in upregulation of lipogenic factors, accumulation of hepatic lipids, and liver steatosis^48^. In the future, careful mechanistic studies on Hh signaling, which account for secondary effects from additional metabolic regulators, will be needed to define the Gli-dependent and -independent contributions towards the governance of lipid homeostasis in the liver.

The catabolic clearance of cellular debris through autophagy is generally thought to benefit longevity; multiple paradigms that extend lifespan are at least partially dependent on increased autophagy in several model organisms^49^. However, in the context of increased mitochondrial permeability, elevated levels of autophagy have the opposite effect on lifespan^13^. SGK-1 regulates mitochondrial permeability transition pore (mPTP) opening; and thus, *rict-1*/*sgk-1* mutants demonstrate a markedly shortened lifespan despite increased levels of autophagy^13^. Consistently, Hedgehog mutants are short-lived but show increased levels of autophagic markers, and their lifespan is modestly extended with knockdown of *hlh-30*/*TFEB*. This supports our hypothesis that Hedgehog signaling positively regulates mTORC2 to modulate its downstream targets, including those regulating autophagy. However, we cannot completely exclude the possibility that Hedgehog works in parallel with mTORC2 to repress autophagy. Negative regulation of autophagy by canonical Hedgehog signaling, requiring the Gli2/Ci transcription factor, has been demonstrated in mammalian cells and *Drosophila*^50^. Contradictorily, both activation and inhibition of non-canonical Hedgehog signaling have been reported to induce autophagy and lipid catabolism in murine hepatocytes via Smo-dependent regulation of AMPK^33^. Complex signaling networks are initiated at the level of Smo, which may stimulate diverse responses through canonical and multiple non-canonical arms of the Hedgehog signaling pathway^51^. This can complicate the interpretation of SAG-induced responses, like autophagy, that are regulated by multiple homeostatic pathways. Our data suggest that non-canonical Hedgehog signaling, through downstream activation of mTORC2 signaling and independent of Gli/TRA-1, inhibits autophagy. Additional studies will be needed to elucidate the mechanisms by which the non-canonical Hedgehog-mTORC2 signaling axis regulates autophagy.

Canonical Hedgehog signaling proceeds through multiple signal transducers, including Smo, and culminates in the activation of the Gli/Ci transcription factor. Primary cilia are required for canonical Hedgehog transcriptional responses through Gli in mammals, but not *D. melanogaster*^52^. *C. elegans* have no known ortholog of Smo, their intestinal cells are not ciliated, and we provide evidence that Gli/TRA-1 is not responsible for the metabolic responses to Hedgehog perturbation, making this system an ideal model to investigate non-canonical Hedgehog signaling. Our findings in *C. elegans* are supported by experiments in mouse hepatocytes, which also lack cilia^53^, suggesting that non-canonical Hedgehog signaling may be conserved from worms to mammals. Hedgehog can act through a Smo-dependent, non-canonical signaling pathway in cholangiocarcinoma and endothelial cells, for example, to modulate cell migration and cytoskeleton remodeling through stimulation of RhoA and Rac1^54–56^. Our results suggest that Smo-dependent, Gli-independent non-canonical Hedgehog signaling may be a common feature of non-ciliated cells; however, a comprehensive study of cell types lacking primary cilia is needed to assess whether this is a widely employed signaling mechanism.

Hedgehog signaling plays a key role in coordinating metabolism through regulation of homeostatic pathways; however, more work is needed to understand the mechanisms of this crosstalk. It’s unclear how Hedgehog signaling engages mTORC2 and p38 MAPK signaling, as well as how p38 signaling restricts lipid accumulation in both mTORC2 and Hedgehog mutants, while only Hedgehog inhibits p38/PMK-1 phosphorylation. It is possible that the two parallel pathways converge on a common set of transcriptional regulators that govern lipid metabolism. In this model, loss of mTORC2 would not alter p38 phosphorylation, but would nonetheless result in misexpression of p38/PMK-1-dependent genes. Our work establishes a new paradigm that explains how developmental pathways may broadly exert control over metabolism and opens new opportunities to investigate the mechanisms that mediate this inter-tissue signaling axis.

## MATERIALS AND METHODS

### C. elegans strains

All *C. elegans* strains were cultured on NGM media seeded with *E. coli* OP50 as previously described^57^. Strains were grown at 20**°**C or 25**°**C as indicated in the figure legends or methods. The strains used in this study are listed in Supplementary Table S1.

### CRISPR/Cas9 genomic editing

Gene editing was performed by microinjection of Cas9::crRNA:tracrRNA complexes (Integrated DNA Technologies) into the *C. elegans* gonad as previously described^58^. All crRNA sequences are listed in Supplementary Table S2. Nonsense or missense mutations were generated using single-stranded oligodeoxynucleotides as homologous recombination (HR) donor molecules, while knock-in of AID or fluorescent protein cassettes was performed using dsDNA HR donors molecules (∼40 bp homology arms), which were generated via PCR using the Q5 DNA Polymerase (New England BioLabs), purified with HighPrep PCR Clean-up beads (MagBio), and then melted and reannealed prior to germline microinjection^58^. Generation of the *grd-3::HA::grd-3::mKate2::3xFLAG* strain was obtained through two rounds of editing. The *grd-4(rhd134)* deletion allele was generated using the *pha-1* co-conversion method, as previously described^59^.

### RNAi experiments

RNAi knockdown of *lin-29*, *grl-21*, *ptr-24*, *hlh-30*, or *tra-1* gene expression was performed by feeding animals *E. coli* HT115(DE3) strains carrying the individual RNAi plasmids (Ahringer RNAi library). The empty vector pL4440 was used as the negative control. Bacterial strains were cultured for ∼16 hrs at 37°C in Luria–Bertani media containing ampicillin (100 μg/ml), concentrated 20-30x by centrifugation, seeded on NGM plates containing 5 mM isopropyl-β-D-thiogalactoside (IPTG) and 100 μg/ml ampicillin, and kept at room temperature overnight to induce dsRNA expression. In most cases, eggs were picked to RNAi plates, allowed to grow for two generations, and then picked to new RNAi plates at the L4 stage where they grew to adulthood for 24 hours before imaging.

### Auxin-inducible degron experiments

Animals were reared on NGM plates for at least two generations before transferring embryos to NGM plates containing 4 mM Naphthaleneacetic Acid (K-NAA, PhytoTech). At the L4 stage, animals were transferred to new plates containing auxin and allowed to develop for an additional 24 hours prior to imaging. Insertion of the AID tags were on a predicted cytoplasmic loop of CHE-14 and the C-terminus of PTR-24.

### Reporter imaging and quantification

The single-copy *rhdSi42[*P*vit-3::mCherry]* vitellogenesis reporter has been previously described^60^. Generally, transgenic animals were grown asynchronously and L4 stage animals were picked to new plates, grown for an additional 24 hours, mounted on a 2% agarose pad with 25 mM levamisole, and imaged on a Nikon SMZ-18 Stereo microscope equipped with a DS-Qi2 monochrome camera. In some cases, L1 synchronized animals were grown for 72 hours at 20**°**C prior to imaging. Using Fiji (v2.14.0)^61^, animals were outlined in the brightfield channel and the mean gray value (fluorescence intensity) was measured in the mCherry channel for each animal. Body size measurements (pixels/worm) were performed simultaneously and were converted to mm^2^ using the known imaging parameters. Data were plotted in Prism 9 as the mean ± SD and a one-way ANOVA with a Bonferroni correction was performed.

Imaging of GRD-3::mKate2 was performed on a Nikon Ti2 widefield microscope equipped with a Hamamatsu ORCA-Fusion BT camera. To differentiate hypodermal from muscle tissue, we microinjected 5 ng/μl P*myo-3::GFP* and 95 ng/μl of 2-Log DNA ladder (New England BioLabs) into the germline of GRD-3::mKate2 animals to generate transgenics expressing a muscle-specific GFP marker.

### Lipid staining experiments

Day 1 adult animals were stained with either Nile Red or Oil Red O to assess total lipid levels. For Oil Red O staining, animals were fixed in 60% isopropanol before staining with 0.3% Oil Red O for 7 hours as previously described^60^. The stained animals were mounted on a 2% agarose pad and imaged with a Nikon SMZ-18 Stereo microscope equipped with a DS-Qi2 monochrome camera. For Nile Red staining, fixed animals (60% isopropanol) were stained for 2 hours with 0.03 mg/mL Nile Red in 40% isopropanol^62^. Subsequently, the animals were washed in M9, mounted on a 2% agarose pad, and immediately imaged with a Nikon Ti2 widefield microscope equipped with a Hamamatsu ORCA-Fusion BT camera. Quantification of Oil Red O and Nile Red was performed in Fiji exactly as previously described^63^. Data were plotted using Prism 9 and a one-way ANOVA with a Bonferroni correction was performed to calculate statistical significance.

### Lifespan assays

Strains for lifespan analysis were maintained without starvation for at least two generations on standard NGM plates seeded with *E. coli* OP50 plates at the appropriate temperature (either 20**°**C and 25**°**C). During the reproductive period, animals were picked to new plates daily to prevent starvation and to keep them separate from their progeny. Approximately 150 animals were assayed for each *C. elegans* strain and lifespan experiments were performed twice with similar results. Data were plotted with Prism 9 and a log-rank test was used to determine statistical significance.

### Quantitative PCR

Day 1 adults (∼5,000 animals reared at 20**°**C) were harvested in M9 buffer, washed three times, and flash frozen. RNA was prepared using Trizol Reagent (Thermo Fisher) and cDNA was synthesized by oligo(dT) priming with the SuperScript III kit (Thermo Fisher) exactly as previously described^3^. The qPCR primer sequences used to query *vit* gene expression have also been previously described^3^. Data from three independent experiments were plotted as the mean fold change relative to wild-type with the standard error of the mean (SEM) using Prism 9.

### Cell culturing

The AML12 hepatocyte cell line was isolated from a 3-month-old male mouse liver (CD1 strain, line MT42) transgenic for human TGFα (CRL-2254, ATCC). AML12 cells were cultured at 37**°**C in 5% CO_2_ in DMEM/Ham’s F12 50/50 media (Fisher Scientific) with 1X Insulin-Transferrin-Selenium (ThermoFisher), 40 ng/mL dexamethasone (Sigma), 10% fetal bovine serum (FBS, #97068-085, VWR), and 1% penicillin-streptomycin (P/S, ThermoFisher). Cells were serum-starved in DMEM/Ham’s F12 50/50 media with 1% P/S and 0.5% FBS for 24 hours before prior to the assay. For drug treatments, human recombinant insulin (ThermoFisher) was added at 100 nM for 30 minutes and Smoothened agonist (SAG, Cayman Chemicals) was added at 0.5 μM for the indicated times.

### siRNA transfections of AML12 cells

AML12 cells were co-transfected with siGLO Green Transfection Indicator and siRNAs targeting *M. musculus* Rictor (final concentration of 50 nM; siGENOME, SMARTPool, Horizon Discovery) using the DharmaFECT reagent (54 μL/10 cm plate) in OPTI-MEM reduced serum media (ThermoFisher) according to the manufacturer’s instructions (Horizon Discovery). A siGLO transfection lacking siRNAs served as the negative control. Media was changed to 0.5% FBS 24 hours after transfection and cells were assayed 48 hours after transfection.

### Western blot analyses

For *C. elegans* samples, 5,000-10,000 synchronized L1 animals were grown to day 1 of adulthood at 20**°**C, harvested in M9 buffer, washed three times, and snap frozen in M9 with 0.01% Tween-20. Whole cell lysates were prepared in RIPA buffer (Cell Signaling Technology) as previously described^60^ and protein concentrations were measured using the DC Protein Assay (Bio-Rad) or the Qubit Protein Broad Rang Assay kit (Thermo Fisher). For AML12 cell lysates, ∼5×10^6^ cells cultured in a 10 cm dish were harvested by scraping in 1X RIPA buffer with 1X HALT Protease and Phosphatase Inhibitor Cocktail (Thermo Fisher) containing 1 mM PMSF and centrifuged (5 min at 13,000 rpm). Protein concentrations in cleared cell lysates were measured using the Qubit Protein Broad Rang Assay kit. Protein samples (∼50 μg) were resolved by SDS-PAGE, transferred to a PVDF membrane, and probed with either anti-FLAG (M2, Sigma, F1804), anti-HA (3F10, Sigma), anti-Actin (ab3280, Abcam), anti-pAkt^S473^ (D9E, Cell Signaling Technology), anti-pAkt^T308^ (D25E6, Cell Signaling Technology), anti-Akt (Cell Signaling Technology), anti-GLI1 (C68H3, Cell Signaling Technology), anti-pFoxO1/3a/4^T24/T32/T28^ (4G6, Cell Signaling Technology), anti-FoxO1 (C29H4, Cell Signaling Technology), anti-Rictor (Cell Signaling Technology), anti-GFP (GF28R, Invitrogen), anti-phospho-p38^T180/Y182^ for *C. elegans* (3D7, Cell Signaling Technology), anti-phospho-p38^T180/Y182^ for mammalian samples (9211, Cell Signaling Technology), anti-p38 for *C. elegans* (gift from Dr. Read Pukkila-Worley), or anti-p38 for mammalian samples (9212, Cell Signaling Technology) antibodies. Experiments were performed at least twice with similar results. Western blot quantification was done using Fiji (v2.14.0) as previously described^64^.

### mRNA sequencing

Total RNA was prepared from wild-type and mutant day 1 adult animals cultured at 20**°**C as described above (see quantitative PCR). Three independent biological replicates were prepared for each condition. For our initial sequencing studies (Figures 1-5), we prepared the mRNA-Seq libraries using 1 μg of total RNA and the TruSeq RNA Library Prep Kit v2 according to the manufacturer’s instructions (Illumina). The samples were sequenced on an Illumina HiSeq 4000 instrument (single-end, 50bp) at the High Throughput Genomic Sequencing Facility at the University of North Carolina at Chapel Hill. For our subsequent study on the p38 pathway (Figure 6), we generated new wild-type and mutant samples in triplicate, isolated the RNA, and the mRNA-Seq libraries were prepared and sequenced (Illumina, paired-end 150) by the Novogene Corporation (Sacramento, CA). Raw sequencing reads were aligned to the *C. elegans* genome (WS260), gene counts were calculated as previously described^3^, and RPKM values and differentially expressed genes (1% FDR) were generated using the DESeq2 software package^65^. Enrichment for differential gene expression in each of the major *C. elegans* tissues was calculated as the number of observed differentially expressed genes in each group relative to the expected number based on random chance. Gene lists for each tissue have been previously described^29^. Gene ontology analyses were performed using the WormCat algorithm^32^. Finally, bar and scatter plots displaying differential expression data (calculated from the DESeq2 RPKM values) were generated using Prism 9. The raw and processed mRNA-Seq data have been deposited in GEO.

## DATA AVAILABILITY STATEMENT

The raw data for the mRNA-Seq expeiments has been deposited in the Gene Expression Omnibus. All additional raw data that support the conclusions described in this article will be made available by the authors, without undue reservation.

## AUTHOR CONTRIBUTIONS

K.R.V., M.A.N., and R.H.D designed the experiments and K.R.V., M.A.N., P.C.B, L.A.H., and R.H.D performed the experiments and interpreted the results. K.R.V. and R.H.D prepared the manuscript.

## FUNDING

This study was supported by the National Institute of General Medical Sciences grants T32GM007092 to K.R.V. and R35GM137985 to R.H.D.

## ACKNOWLEDGEMENTS

The Caenorhabditis Genetics Center provided some of the strains used in this study, which is supported by the NIH Office of Research Infrastructure Programs (P40 OD010440). We thank XXX.

## CONFLICT OF INTEREST

The authors declare no competing interests.

## Notes

### Competing Interest Statement

The authors have declared no competing interest.

