## Supplemental Material for "Non-cell-autonomous regulation of mTORC2 by Hedgehog signaling maintains lipid homeostasis"

**Contents** **Page**

1. Supplemental Tables S1-S2 2
2. Supplemental Figures S1-S4 5
3. References 9

| **Strain** | **Genotype** | **Reference** |
|---|---|---|
| N2 | Wild-type | (Brenner, 1974) |
| BL3466 | *inIs173[Pvit-2::NLS-GFP] II* | (MacMorris et al., 2003) |
| CB3698 | *che-14(e1960) I* | (Perkins et al., 1986) |
| DA2123 | *adIs2122[Plgg-1::GFP::lgg-1 + rol-6(su1006)]* | (Nakae et al., 2010) |
| DLS433 | *rhdSi39[Pnhr-73::mCherry::his-58::SL2::lin-29c + cb-unc-119(+)] II* | This study |
| DLS439 | *rhdSi35[Pvha-6::mCherry::his-58::SL2::lin-29c + cb-unc-119(+)] lin-29(n333) II* | This study |
| DLS487 | *che-14(ok193) I; grd-3(ok2778) IV; grd-4(rhd134) X* | This study |
| DLS488 | *che-14(e1960) I; grd-3(ok2778) IV; grd-4(rhd134) X* | This study |
| DLS490 | *rict-1(mg360) II* | (Jones et al., 2009) |
| DLS493 | *grd-3(ok2778) IV; grd-4(rhd134) X* | This study |
| DLS506 | *inIs173[Pvit-2::NLS-GFP] II; grd-3(ok2778) IV; grd-4(rhd134) X* | This study |
| DLS507 | *che-14(ok193) I; inIs173[Pvit-2::NLS-GFP] II; grd-3(ok2778) IV; grd-4(rhd134) X* | This study |
| DLS524 | *inIs173[Pvit-2::NLS-GFP] II; grd-3(ok2778) IV; grd-4(rhd134) sgk-1(ft15) X* | This study |
| DLS525 | *inIs173[Pvit-2::NLS-GFP] II; grd-3(ok2778) IV; akt-1(mg144) V; grd-4(rhd134) X* | This study |
| DLS537 | *rhdSi42[Pvit-3::mCherry::unc-54 3'UTR + cb-unc-119(+)] II* | (Torzone et al., 2023) |
| DLS539 | *rhdSi42[Pvit-3::mCherry::unc-54 3'UTR + cb-unc-119(+)] alxIs9[Pvha-6::SID-1::SL2::GFP] II; sid-1(qt9) V* | (Torzone et al., 2023) |
| DLS540 | *rhdSi42[Pvit-3::mCherry::unc-54 3'UTR + cb-unc-119(+)] II; grd-3(ok2778) IV; grd-4(rhd134) X* | This study |
| DLS541 | *rhdSi42[Pvit-3::mCherry::unc-54 3'UTR + cb-unc-119(+)] II; grd-3(ok2778) IV; grd-4(rhd134) sgk-1(ft15) X* | This study |
| DLS551 | *rhdSi42[Pvit-3::mCherry::unc-54 3'UTR + cb-unc-119(+)] II; grd-3(ok2778) IV; akt-1(mg144) V; grd-4(rhd134) X* | This study |
| DLS594 | *daf-16(ot821[daf-16::mKate2::3xFLAG]) I; pqm-1(rhd187[BioTag::GFP::TEV::3xFLAG::pqm-1]) II* | This study |
| DLS622 | *reSi5[Pges-1::TIR1::F2A::mTagBFP2::NLS::AID::tbb-2 3'UTR] I; rhdSi42[Pvit-3::mCherry::unc-54 3'UTR + cb-unc-119(+)] II* | (Torzone et al., 2023) |
| DLS710 | *reSi5[Pges-1::TIR1::F2A::mTagBFP2::NLS::AID::tbb-2 3'UTR] I; rhdSi42[Pvit-3::mCherry::unc-54 3'UTR + cb-unc-119(+)] II; grd-3(ok2278) IV; grd-4(rhd134) X* | This study |
| DLS754 | *alxIs9[Pvha-6::sid-1::SL2::GFP] II; sid-1(qt9) V; sgk-1(ok538) X* | This study |
| DLS788 | *grd-3(rhd252[grd-3(1-32aa)::HA::grd-3(33-181aa)::mKate2::3xFLAG]) IV* | This study |
| DLS797 | *wrdSi23[Peft-3::TIR1::F2A::BFP::AID*::NLS::tbb-2 3'UTR] che-14(rhd259[che-14::3xFLAG::AID::che-14]) I; rhdSi42[Pvit-3::mCherry::unc-54 3'UTR + cb-unc-119(+)] II* | This study |
| DLS798 | *ptr-21(rhd260[S122*, M123I, R125R]) I; rhdSi42[Pvit-3::mCherry::unc-54 3'UTR + cb-unc-119(+)] II* | This study |

| | | |
|---|---|---|
| DLS799 | *ptr-21(rhd261[S122*, M123I, R125R]) I; rhdSi42[Pvit-3::mCherry::unc-54 3'UTR + cb-unc-119(+)] II* | This study |
| DLS800 | *ptr-21(rhd262[M123fs]) I; rhdSi42[Pvit-3::mCherry::unc-54 3'UTR + cb-unc-119(+)] II* | This study |
| DLS801 | *ptr-21(rhd263[M123fs]) I; rhdSi42[Pvit-3::mCherry::unc-54 3'UTR + cb-unc-119(+)] II* | This study |
| DLS828 | *reSi1[Pcol-10::TIR1::F2A::mTagBFP2::AID*::NLS::tbb-2 3'UTR] che-14(rhd259[che-14::mNG::3xFLAG::AID]) I; rhdsi42[Pvit-3::mCherry::unc-54 3'UTR + cb-unc-119(+)] II* | This study |
| DLS829 | *reSi5[Pges-1::TIR1::F2A::mTagBFP2::NLS::AID::tbb-2 3'UTR] che-14(rhd259[che-14::mNG::3xFLAG::AID]) I; rhdsi42[Pvit-3::mCherry::unc-54 3'UTR + cb-unc-119(+)] II* | This study |
| DLS866 | *che-14(rhd259[che-14::3xFLAG::AID::che-14]) I; rhdSi42[Pvit-3::mCherry::unc-54 3'UTR + cb-unc-119(+)] II* | This study |
| DLS938 | *rict-1(mg360) II; tir-1(qd4) III* | This study |
| DLS943 | *che-14(ok193) I; tir-1(qd4) III; grd-3(ok2778) IV; grd-4(rhd134) X* | This study |
| DLS945 | *rict-1(mg360) II; sqIs19[Phlh-30::HLH-30::GFP + rol-6(su1006)]* | This study |
| DLS956 | *che-14(ok193) I; rhdSi42[Pvit-3::mCherry::unc-54 3'UTR + cb-unc-119(+)] II; grd-3(ok2778) IV; grd-4(rhd134) X* | This study |
| DLS957 | *che-14(ok193) daf-16(ot821[daf-16::mKate2::3xFLAG]) I; pqm-1(rhd187[BioTag::GFP::TEV::3xFLAG::pqm-1]) II; grd-3(ok2778) IV; grd-4(rhd134) X* | This study |
| DLS959 | *che-14(ok193) I; grd-3(ok2778) IV; grd-4(rhd134) X; adIs2122[lgg-1p::GFP::lgg-1 + rol-6(su1006)]* | This study |
| DLS963 | *grd-3(ok2778) IV; grd-4(rhd134) X; sqIs19[Phlh-30::HLH-30::GFP + rol-6(su1006)]* | This study |
| DLS970 | *reSi5[Pges-1::TIR1::F2A::mTagBFP2::NLS::AID::tbb-2 3'UTR] I; rhdSi42[Pvit-3::mCherry::unc-54 3'UTR + cb-unc-119(+)] II; grd-3(ok2278) IV; grd-4(rhd134) ptr-24(rhd317[ptr-24::2xminiIAA7::3xHA] X* | This study |
| MAH235 | *sqIs19[Phlh-30::HLH-30::GFP + rol-6(su1006)]* | (Lapierre et al., 2013) |
| MGH171 | *sid-1(qt9) V; alxIs9[Pvha-6::SID-1::SL2::GFP]* | (Melo & Ruvkun, 2012) |
| ML514 | *che-14(ok193) I* | (Michaux et al., 2000) |
| MT333 | *lin-29(n333) II* | (Ambros & Horvitz, 1984) |
| RB2103 | *grd-3(ok2778) IV* | (*C. elegans* Deletion Mutant Consortium, 2012) |
| RPW43 | *nsy-1(ums8) II; agIs44[pF08G5.6::GFP::unc-54(3'UTR) + Pmyo-2::mCherry]* | (Cheesman et al., 2016) |
| ZD101 | *tir-1(qd4) III* | (Shivers et al., 2009) |

**Supplementary Table S1. *C. elegans* strains in this study.** Strain names, genotypes, and references are listed for strains used in this study.

| Target gene | Location in gene (crRNA guide number) | crRNA sequence | Alleles | Genomic edit |
|---|---|---|---|---|
| *grd-3* | N-terminal (rhd37); internal (rhd17) | rhd37: GAUCGUCAGCUUGCCAAGCC<br>rhd17: GAAUCCUCCAGCAAGGCCGC | *rhd252* | HA tag (rhd37)<br><br>mKate2::3xFLAG (rhd17) |
| *pqm-1* | N-terminal (rhd8) | CAUUAUUCAAAAACGACAUU | *rhd187* | BioTag::GFP:: 3xFLAG |
| *che-14* | Internal (rhd35) | CACGAAGAGCUGCUGCCAAU | *rhd259* | 3xFLAG::AID |
| *ptr-21* | Internal (rhd22) | AAGACGGAGGAUCCAUGCAU | *rhd260, rhd261, rhd262, rhd263* | *rhd260*, S122*<br>*rhd261*, S122*<br>*rhd262*, M123fs, 138bp insertion<br>*rhd263*, M123fs, 92bp insertion |
| *ptr-24* | C-terminal (rhd28) | AGAAAUUAAUUAAGAUUGAG | *rhd317* | 2xmIAA7::3xHA |

**Supplementary Table S2. The crRNAs used in this study.** List of all crRNA guides, target genes, ribonucleotide sequences, alleles, and a description of the genomic edits in this study.

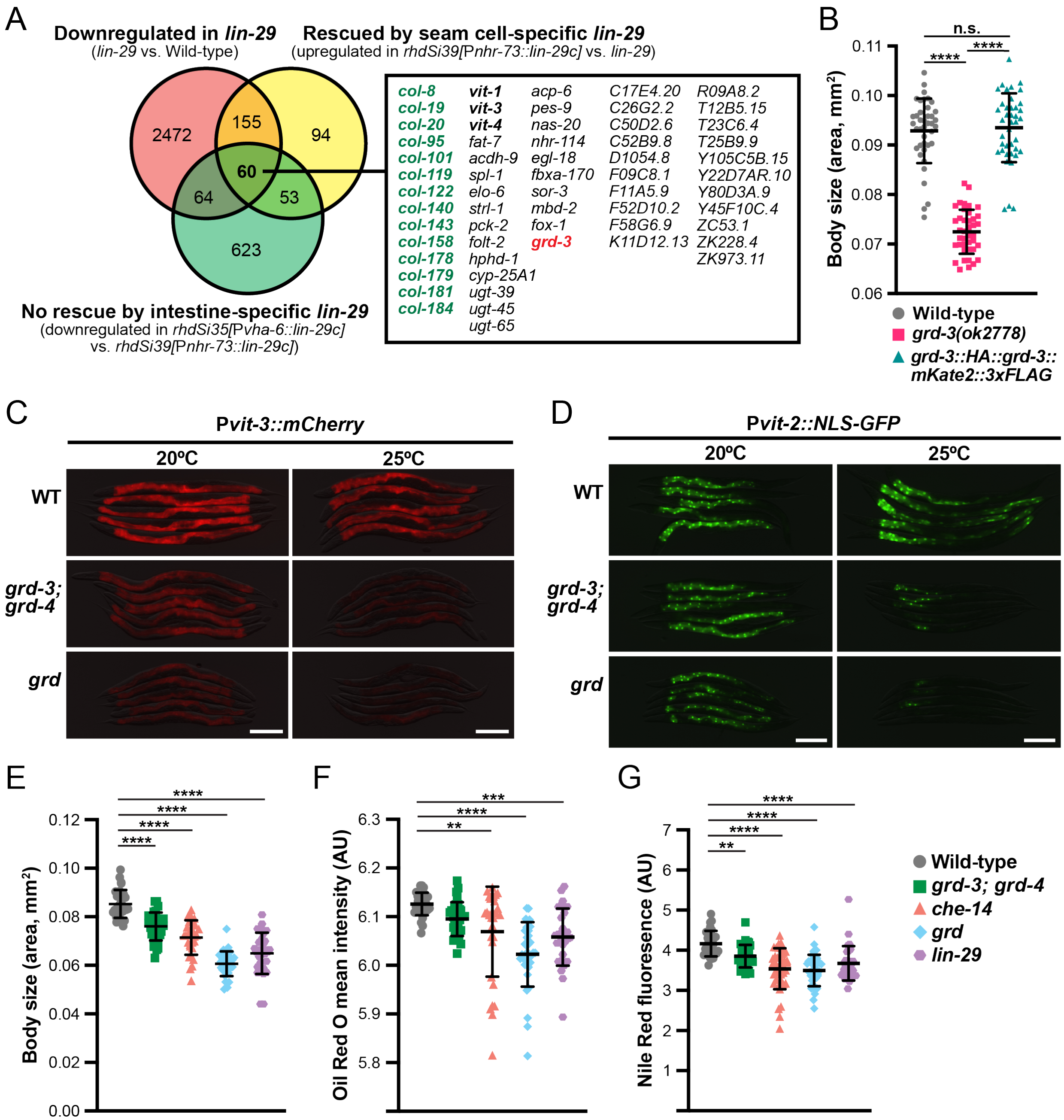


**Figure S1. Identification of Hedgehog signaling as a master regulator of lipid metabolism.** (A) A Venn diagram of genes down-regulated in the *lin-29(n333)* mutant, genes up-regulated upon seam cell-specific rescue of *lin-29* (relative to the *lin-29(n333)* mutant), and genes whose expression was not altered by intestine-specific rescue of *lin-29* (non-specific regulation) . This analysis resulted in the identification of 60 genes, including multiple *vit* genes (black), several adult-specific collagens (green), and *grd-3* (red). (B) A body size analysis of wild-type, *grd-3(ok2778)*, and *grd-3(rhd252[grd-3(1-32aa)::HA::grd-3(33-181aa)::mKate2::3xFLAG])* animals (mean ± SD, ****, *P*<0.0001, n.s., not significant, one-way ANOVA), demonstrating that the epitope tags in GRD-3 do not disrupt protein function. Representative (C) P*vit-3::mCherry* and (D) P*vit-2::NLS-GFP* flourescence images of wild-type, *grd-3(ok2778); grd-4(rhd134)* double mutant, and *grd-3(ok2778); grd-4(rhd134); che-14(ok193) (grd)* triple mutant animals reared at either 20°C or 25°C (scale bars, 200 μm). (E) Body size measurements and quantification of (F) Oil Red O and (G) Nile Red staining of day 1 adult wild-type, *lin-29(n333)*, and Hh mutant animals reared at 20°C. All data are reported as the mean ± SD (****, *P*<0.0001, ***, *P*<0.001, **, *P*<0.01, one-way ANOVA).

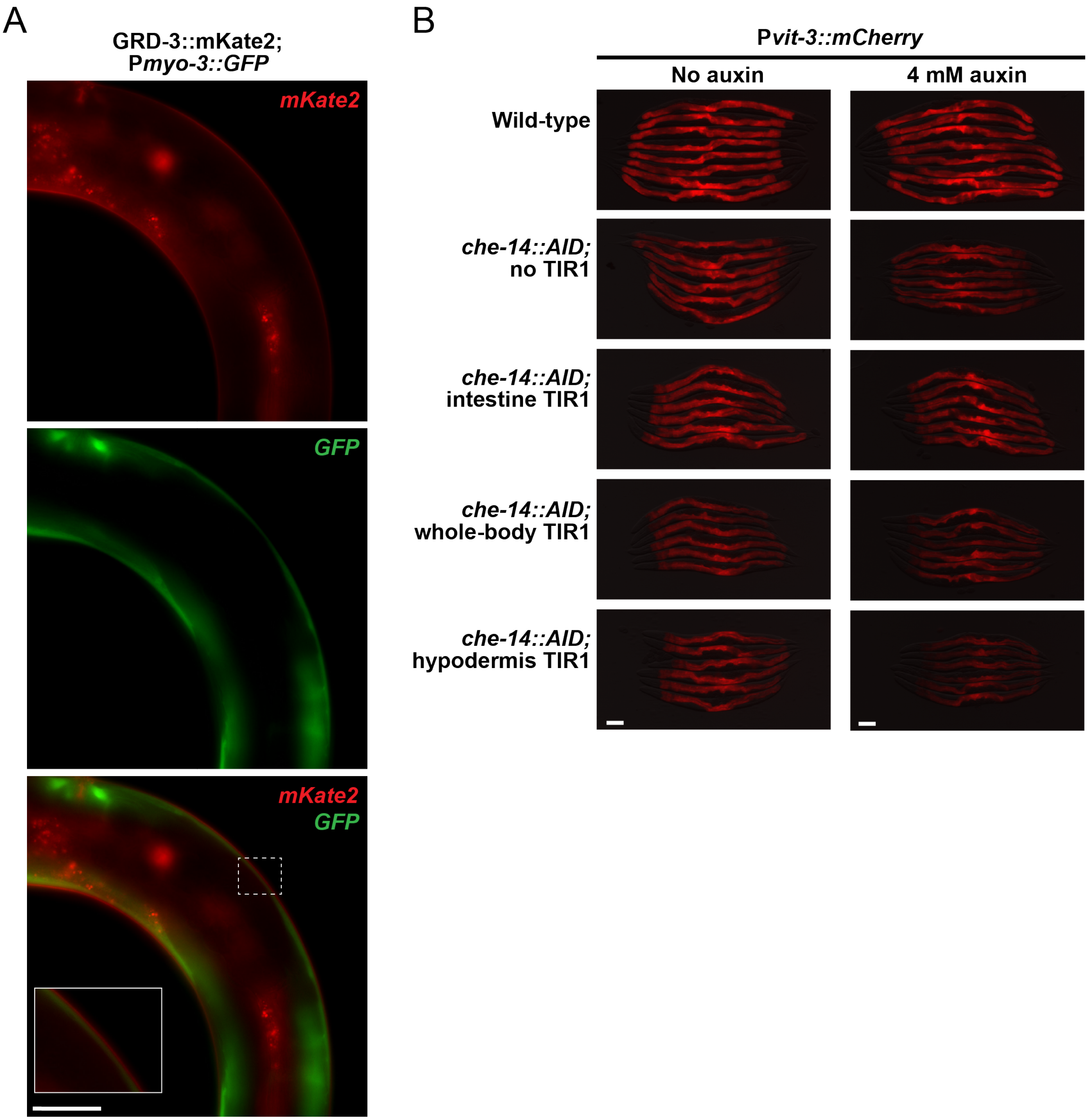


**Figure S2. The Hh ligands are synthesized and dispatched from hypodermal tissue.** (A) Representative flourescence images of an animal expressing the GRD-3::mKate2 protein (red, top) and a P*myo-3::GFP* muscle marker (green, middle). The overlaid image is also displayed (bottom; scale bar, 50 µm), demonstrating that GRD-3::mKate2 is not produced in the muscle tissue. White dashed box is cropped and zoomed in the bottom left. (B) Representative P*vit-3::mCherry* flourescence images of *che-14::AID* animals cultured in the absence or presence of 4 mM auxin at 20°C (scale bars, 50 µm). Depletion of CHE-14 in hypodermal cells reduces P*vit-3::mCherry* reporter expression to largest degree.

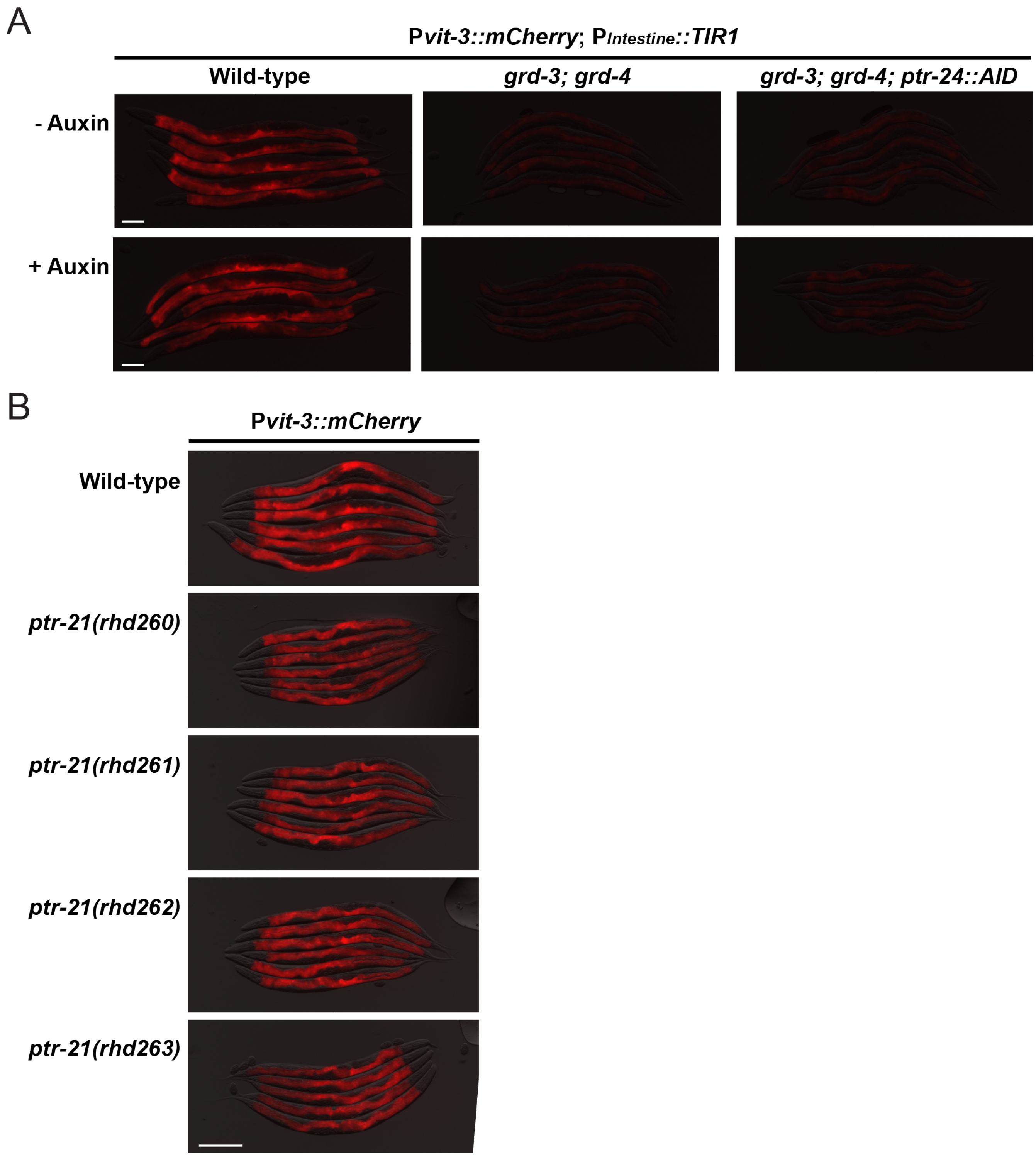


**Figure S3. Identification of a candidate Patched-related protein that is partially required for vitellogenin synthesis.** (A) Representative P*vit-3::mCherry* flourescence images of wild-type, *grd-3(ok2778); grd-4(rhd134)*, or *grd-3(ok2778); grd-4(rhd134); ptr-24::AID* animals reared in the absence or presence of 4 mM auxin (25°C; scale bar, 100 µm). Depletion of PTR-24 in intestinal cells does not restore P*vit-3::mCherry* reporter expression in the Hh ligand mutant. (B) Representative P*vit-3::mCherry* flourescence images of wild-type and four different *ptr-21* loss-of-function mutants (20°C; scale bar, 200 µm). Loss of *ptr-21* causes a modest decrease in P*vit-3::mCherry* reporter expression.

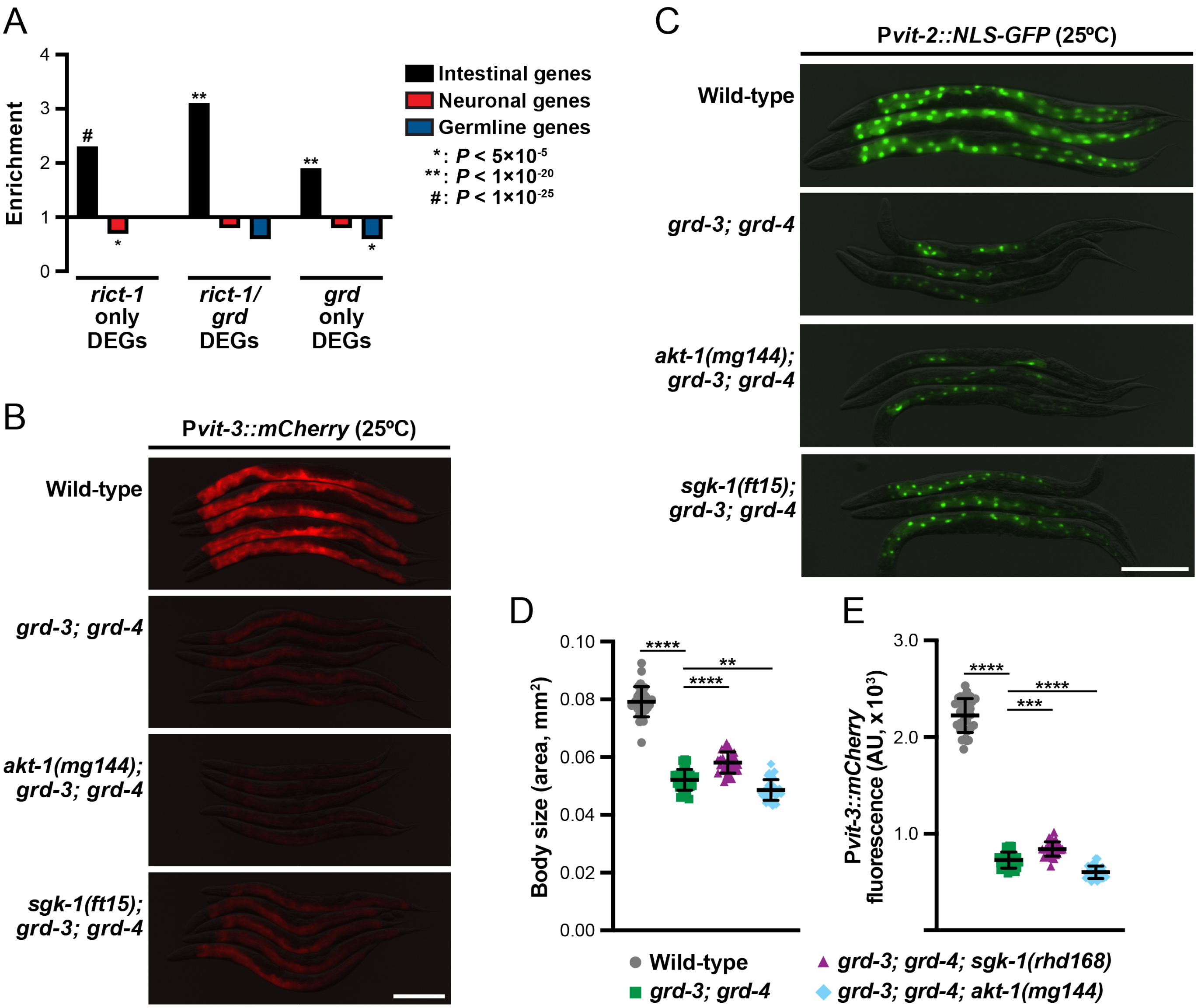


**Figure S4. Intestinal mTORC2 signaling acts genetically downstream of Hh.** (A) Enrichment (bars >1) or depletion (bars <1) of differentially expressed genes within the indicated tissues, indicating that *rict-1/grd* co-regulated genes are enriched in the intestine (relative to random chance). Hypergeometric *P* values are reported. Representative (B) P*vit-3::mCherry* and (C) P*vit-2::NLS-GFP* flourescence images of day 1 adult wild-type, *grd-3(ok2778); grd-4(rhd134)*, *grd-3(ok2778); grd-4(rhd134); akt-1(mg144),* and *grd-3(ok2778); grd-4(rhd134); sgk-1(ft15)* animals reared at 25°C (scale bars, 200 µm). Quantification of (D) body size and (E) P*vit-3::mCherry* flourescence of day 1 adult wild-type and Hh mutant animals cultured at 25°C (mean ± SD, ****, $P$<0.0001, ***, $P$<0.001, **, $P$<0.01, one-way ANOVA). The *ft15* and *rhd168 sgk-1* alleles are gain-of-function mutations.
